# Phylogenomic and genomic analysis reveals unique and shared genetic signatures of *Mycobacterium kansasii* complex species

**DOI:** 10.1101/2024.02.14.579889

**Authors:** Edson Machado, Sidra Vasconcellos, Lia Gomes, Marcos Catanho, Jesus Ramos, Luciana de Carvalho, Telma Goldenberg, Margareth Dalcolmo, Paulo Redner, Paulo Caldas, Carlos Campos, Maria Cristina Lourenço, Elena Lasunskaia, Vinicius Mussi, Lizania Spinassé, Solange Vinhas, Leen Rigouts, Sari Cogneau, Pim de Rijk, Christian Utpatel, Jarmila Kaustova, Tridia van der Laan, Han de Neeling, Nalin Rastogi, Klavdia Levina, Marge Kütt, Igor Mokrousov, Viacheslav Zhuravlev, Ndivhu Makhado, Manca Žolnir-Dovč, Vera Jankovic, Jacobus de Waard, Maria Sisco, Dick van Soolingen, Stefan Niemann, Bouke C. de Jong, Conor J. Meehan, Philip Suffys

## Abstract

Species belonging to the *Mycobacterium kansasii* complex (MKC) are frequently isolated from humans and the environment and can cause serious diseases. The most common MKC infections are caused by the species *M. kansasii* (*stricto sensu*), leading to tuberculosis-like disease. However, a broad spectrum of virulence, antimicrobial resistance and pathogenicity of these non-tuberculous mycobacteria (NTM) are observed across the MKC. Many genomic aspects of the MKC that relate to these broad phenotypes are not well elucidated. Here, we performed genomic analyses from a collection of 665 MKC strains, isolated from environmental, animal and human sources. We inferred the MKC pangenome, mobilome, resistome, virulome and defense systems and show that the MKC species harbors unique and shared genomic signatures. High frequency of presence of prophages and different types of defense systems was observed. We found that the *M. kansasii* species splits into four lineages, of which three are lowly represented and mainly in Brazil, while one lineage is dominant and globally spread. Moreover, we show that four sub-lineages of this most distributed *M. kansasii* lineage emerged during the 20^th^ century. Further analysis of the *M. kansasii* genomes revealed almost 300 regions of difference contributing to genomic diversity, as well as fixed mutations that may explain the *M. kansasii*’s increased virulence and drug resistance.

**Repositories:** BioProject PRJNA1048499.

**Impact statement:** *Mycobacterium kansasii* complex (MKC) is a group of closely related non-tuberculous mycobacteria species, recognized as a significant source of human infection. Species belonging to the MKC may present a broad spectrum of virulence, antimicrobial resistance and pathogenicity and there is a lack of knowledge about their genomic content related to these broad phenotypes. We have provided whole genomic sequencing DNA for 342 MKC isolates and, together with public data, investigated the MKC pangenome, mobilome, resistome, virulome and defense systems and show unique and shared genetic signatures within MKC species. Furthermore, with phylogenomic and bayesian population analysis, we inferred the distribution and emergence of the Mycobacterium kansasii species lineages and sub-lineages. This study has considerably expanded the MKC available data, by providing genomic sequences of isolates from countries and global regions with unknown or poorly MKC data until now.

**Data summary:** NGS data generated in the study are available in the NCBI Sequence Read Archive (SRA) repository under the accession number PRJNA1048499 and the accession numbers for all data sets used are provided in Supplementary Table 1.

## Introduction

The *Mycobacterium kansasii* complex (MKC) is a group of closely related non-tuberculous *Mycobacterium* (NTM) species. Currently, the MKC includes *Mycobacterium gastri* and six *Mycobacterium kansasii* subtypes that were recently redefined as distinct species based on genomic taxonomy and phylogenomic analyses^1,2^. Thus, the former *M. kansasii* subtypes I to VI were renamed as *Mycobacterium kansasii* (*stricto sensu*)*, Mycobacterium persicum, Mycobacterium pseudokansasii, Mycobacterium ostraviense, Mycobacterium innocens,* and *Mycobacterium attenuatum*.

In the clinical niche, *M. kansasii* is the most common MKC species isolated, followed by *M. persicum*, which is mainly associated with individuals infected by human immunodeficiency virus (HIV), whereas another MKC species members remain predominantly colonizers and rarely associated with disease^3^. Similar to some other NTM species, strains of the MKC can also be isolated from animal and environmental sources. Besides the pathogenicity and isolation source diversity, MKC species display heterogeneity regarding phenotypic aspects such as virulence, colony morphology and drug resistance. For example, regarded to virulence, pulmonary disease underlying respiratory comorbidities or immunosuppression is the main clinical manifestation of human infections caused by MKC, although the occurrence of extrapulmonary diseases is also observed^4,5^. Furthermore, lung disease caused by *M. kansasii* can result in a broad pathology spectrum^6^.

At the genomic level, the basis of the MKC pathogenicity and phenotypic traits has been explored. Plasmids are estimated to be present in around 30% of MKC isolates, although with no observed correlation with pathogenicity^7^. Conversely, an association of pathogenicity with genetic recombination has been observed, mediated by distributive conjugal transfer, also with possible correlation to MKC speciation events^7,8^. Regarding virulence, whereas *M. kansasii* contains a type-VII secretion system ESX-1 locus with all corresponding orthologs of the *M. tuberculosis* genes, some MKC species do not have an intact conservation of this locus^7,9^. Furthermore, nearly three hundred shared orthologs between *M. kansasii* and *M. tuberculosis* are predicted to encode potential virulence factors^6^ (VFs), but the conservation of these VFs in another MKC species has not been explored.

Concerning the resistance of members of the MKC to antimicrobial agents, estimates are variable, but some recent studies conducted in China^10^ and Brazil^11^ estimated a high percentage (∼70%) of *M. kansasii* isolates resistant to ethambutol and ciprofloxacin, while between 10% and 20% of the isolates were resistant to rifampicin. However, in *M. kansasii*, no clear relation has been observed between mutations in particular genes and drug resistance as observed in *M. tuberculosis*^12^.

Due to the higher interest in isolating MKC from clinical rather that from environmental sources, and the strong association of *M. kansasii* with pulmonary disease, much more genomic sequencing data are available from the latter species, with the majority of data coming from Europe, China, and Australia leaving knowledge of other geographical areas lacking^8,10^. From *M. kansasii* genomic data, it was estimated that some lineages might be globally spread^8^, while genomes obtained from clinical isolates in Brazil and China^10,13^ suggested at least four genetically diverse groups.

Even though some questions about the MKC pathogenicity, virulence, drug resistance, genetic diversity, and epidemiology have been addressed by analyzing genomic data^1,2,6–10,12–14^, several aspects about the MKC species genomic characteristics and epidemiology remains unexplored. For instance, i) in addition to plasmids, the prevalence and characteristics of mobile genetic elements in MKC genomes, such as prophages; ii) the possible plasmids or prophages contribution to virulence or drug resistance; iii) the MKC pangenome structure and diversity; iv) the MKC viral defense mechanisms repertoire; v) the potential *M. kansasii* lineages and their association with certain geographical areas or niches; and vi) transmission mechanisms of the different MKC species.

We have therefore sequenced the genomes of nearly four hundred MKC isolates and together with publicly available data, performed several *in silico* analyses to further address the opened questions mentioned above. In this study we provide a global phylogeny and a genomic panorama of the MKC, that provides new knowledge about the MKC mobilome, resistome, virulome, pangenome, viral defense systems, population structure and genetic diversity.

## Results

### Global distribution and phylogeny of the *Mycobacterium kansasii* complex

To investigate the global distribution and genomic repertoire of the *M*. *kansasii* complex, we gathered whole genome sequencing (WGS) data of 676 strains, of which 342 were collected and sequenced in this study and 334 were obtained from public repositories with assembled genomes or available DNA short sequencing data (Supplementary Table 1). After applying quality thresholds, 665 genome assemblies of strains from 28 countries and five continents remained for genomic and phylogenomic analyses. The isolation source distribution of the genomes is as follows: 538 isolated from humans, 76 from the environment, six from animals and 45 of unknown origin. Information on the colony morphology was available for 142 isolates (112 smooth and 30 rough) and HIV testing results were provided for patients associated with 44 isolates, including 24 positive and 20 negative (Supplementary Table 2, Figure 1).

**Figure 1.**
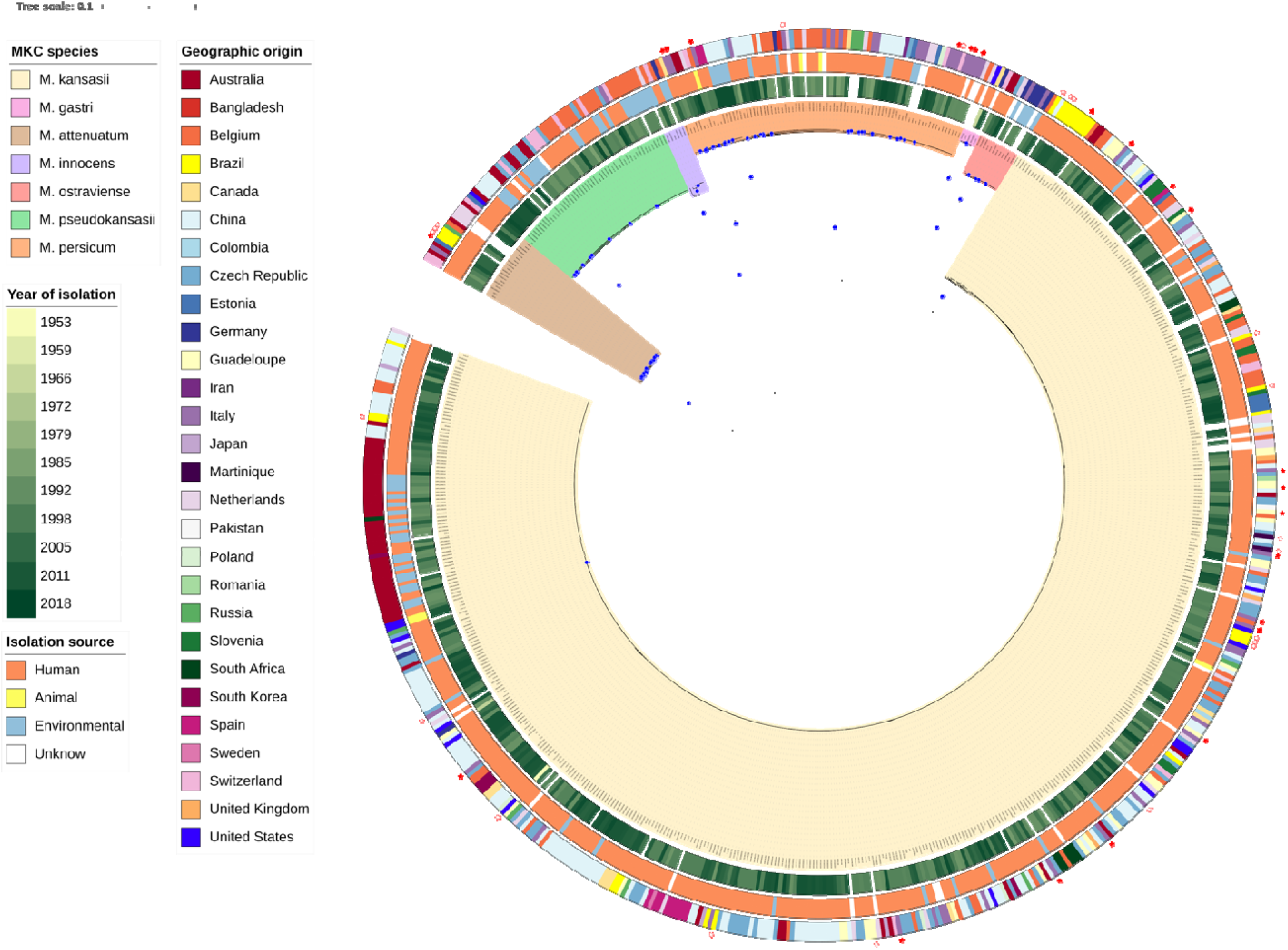
Midpoint rooted maximum likelihood phylogenetic tree of 665 M. kansasii complex genomes. Blue dots indicate nodes with bootstrap support above 75% with shading of branche and nodes indicating the seven MKC species. The rings, from inside towards outside, designate: (i) the year of isolation (if available), (ii) the isolation source of the strains classified as either environmental, animal, human or unknown; (iii) the country of isolation. The outer red stars indicate clinical isolates with patient HIV information, bring positive (solid) or negative (open).

The genome sizes ranged from 5.63LMb to 6.90LMb and GC content was between 65.59 and 66.38%. From the data collected in this study, we performed long-read sequencing technology for three genomes, resulting in fully closed genomes for two samples (KAN-960446 and KAN-130495), each presenting two replicons (a plasmid and a chromosome sequences), and one nearly fully closed genome (sample KAN-C06235) that was composed by nine contigs, including a complete plasmid sequence.

### Prevalence and distribution of plasmids among MKC

By applying and combining three *in silico* strategies, we were able to identify 346 contigs classified as putative plasmids (Supplementary Table 3) within 244 MKC genomes (∼37% of 665). Most of these genomes (presented only a single contig classified as deriving from a plasmid (n=169; 69%), while the others contained two to five contigs per genome classified as plasmid-derived. The length of the contigs predicted as plasmid-derived ranged from 5,423 to 237,122 base pairs with a mean and median of 39.6 and 20.1 kb, respectively. Around 47% (163/348) of the putative plasmids encoded one (n=162) or two (n=1) tRNA genes. Five MKC species had plasmid-derived contigs with the following distribution: *M. kansasii* (176/492, 35.7%), *M. persicum* (21/81, 25.9%)*, M. pseudokansasii* (33/50, 66%)*, M. ostraviense* (1/12, 9%), and *M. attenuatum* (14/19, 73.7%); *M. innocens* and *M. gastri* contained no predicted plasmids (*p*L<L0.05, Fisher’s exact test).

Among the 346 plasmid-derived contigs, 283 had no similarity with reference plasmid sequences, whereas 63 were matched via BLASTn. From the hits returned, 51 were similar to described MKC plasmid sequences while 12 were similar to plasmid sequences from other *Mycobacterium* species (*M. intracellulare subsp. chimaera*, n=4; *M. kubicae*, n=2; *M. aubagnense*, n=6). Based on marker genes, at least one conjugative (122/244), mobilizable (161/244) or replicative (130/244) mechanism was identified in each of the 244 genomes with predicted plasmids (Supplementary Table 3, Supplementary Figure 1).

### Prophages diversity and characterization across MKC

A total of 786 prophages were predicted to be distributed across 527 MKC genomes (∼79.3% of 665) (Supplementary Table 4), with one to six prophages per genome. Prophage lengths ranged from 4.5 Kb to 100.2 Kb, and 109 prophages encoded one (n=70), two (n=6), four (n=32) or 21 (n=1) tRNA genes. Considering all MKC species, prophages were not detected in *M. gastri,* while prophage distribution across the other MKC species was as follows: *M. kansasii* (399/492, 81.1%), *M. persicum* (73/81, 91.1%)*, M. pseudokansasii* (36/50, 72%)*, M. ostraviense* (9/12, 75%)*, M. innocens* (3/6, 50%), and *M. attenuatum* (7/19, 36.9%) (*p*L<L0.05, Fisher’s exact test). Although MKC prophages were grouped into 59 clusters, distinct from the 37 mycobacteriophages clusters found in the PhageDB, the employed network based on the comparison and clustering process of prophages sequences, as well as their predicted genes, revealed a relation between most MKC prophages and other known mycobacteriophages, as evidenced by the edges connections indicating their shared genes (Supplementary Table 5, Figure 2). Around 90% of prophages (710/786) were grouped into 20 clusters with at least five similar prophages, and the biggest cluster encompassing 335 prophages specific to *M. kansasii* (Supplementary Figure 2). Regarding the relationship between prophage clusters and host-species, 42 clusters were species-specific (found in only one species), whereas 17 were diverse, being shared by two to five different species. No prophage cluster contained all MKC species.

**Figure 2.**
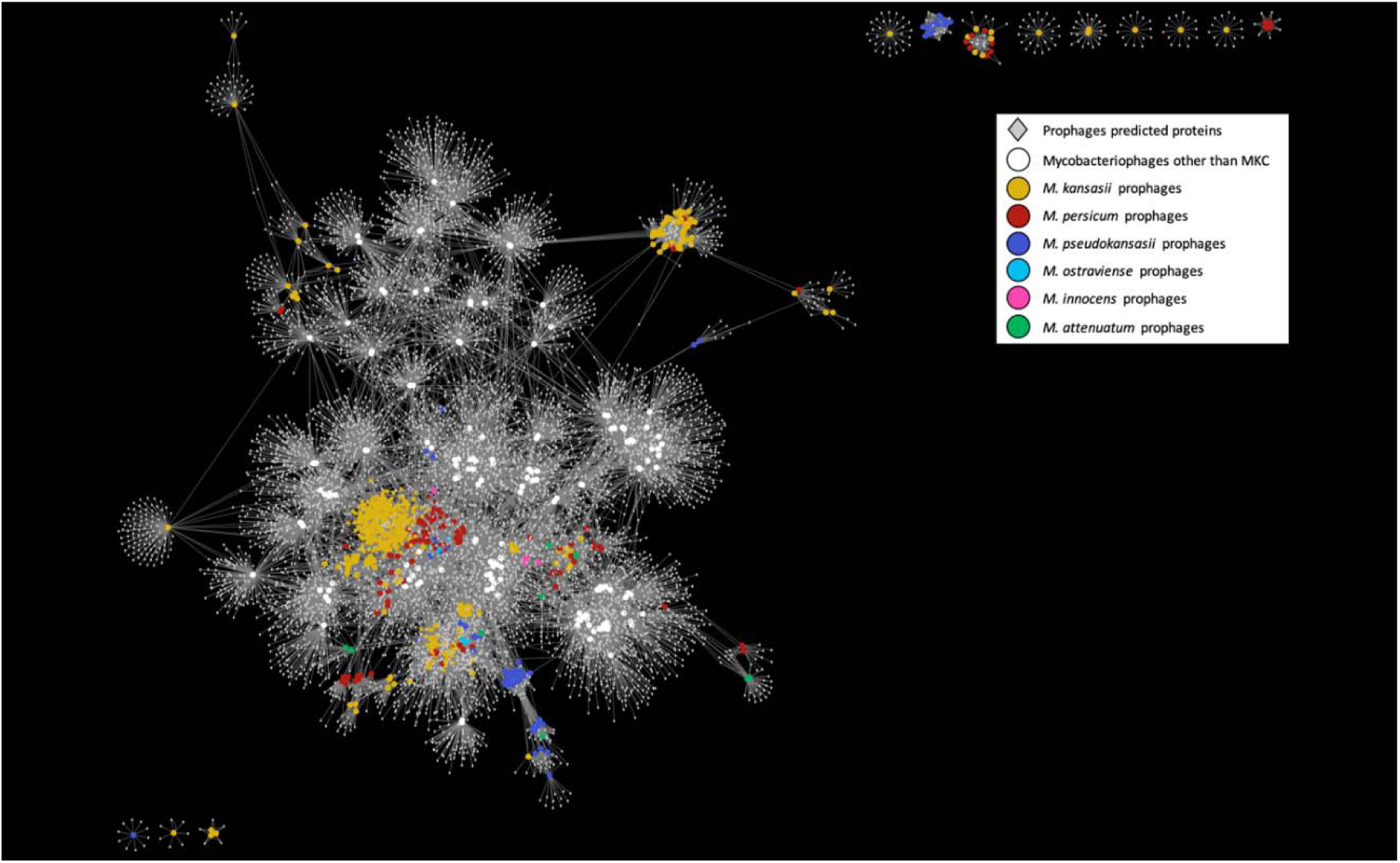
The bipartite network of the MKC prophages and their gene content. The predicted genes of the prophages are show as grey diamonds, whereas prophages genomes as colored circles. Genomes circles are colored by MKC species or Mycobacteriophages genomes obtained from hosts other than MKC species. Genomes are connected to their predicted genes by edges.

### MKC phage types

Among 18 MKC strains, we had information on susceptibility to 11 phage types, as outlined in Supplementary Table 19. Phage type was characterized for 14 isolates of *M. kansasii*, two being phage type 2, 3, 7, 9, 10 and 13 each; a single strain of *M. kansasii* being phage type 6 and 8. Two strains of *M. attenuatum* were types: one was phage type 1 and the other phage type 6. Finally, a strain of *M. ostraviense* and *M. persicum* were of phage type 4 and 14, respectively.

### The MKC pangenome is open and has a highly diverse and species-specific accessory genome

Due to the diverse nature of the complex, we investigated whether different identity cut-offs could substantially impact the MKC pangenome prediction. We generated pangenomes applying 70%, 80% and 90% BLASTp identity cut-offs. For all the three runs performed, the pangenome resulted in a large amount of shell and cloud genes, suggesting a highly diverse accessory genome for the complex. The number of new genes kept increasing regardless of the addition of new genomes, likely characterizing an open pangenome as the increment curve did not achieve a plateau (Supplementary Figure 3).

Intriguingly, with 70% identity cut-off, the resultant core genome (strict and soft) slightly decreased compared with 80% identity. Despite this not expected Roary’s results for the core genome at lower cut-offs, by comparing the pangenome matrices for each cut-off performed, it is clear that each MKC species is characterized by distinct accessory gene content (Supplementary Figure 4).

### The MKC virulome is species specific

Among the 287 genes we detected that were encoding for potential virulence factors, 123 were present in all 665 samples, regardless of the species, while 164 genes were found between 39.9% (265/665) and 99.9% (664/665) of the samples (Supplementary Table 6). The mean and median values of virulence factor count identified per MKC species was as follows: *M. kansasii* (281/283), *M. persicum* (259/260), *M. ostraviense* (251/252), *M. innocens* (247/247), *M. pseudokansasii* (246/248), *M. gastri* (244/244), *M. attenuatum* (229/229) (Supplementary Figure 5). A closer look at the 63 genes belonging to the type VII secretion system (T7SS) that has been described as playing a crucial role in mycobacterial physiology and virulence, showed that only 18 were found in all samples (Supplementary Figure 6). For VFs other than genes of the T7SS, 119 where not found in all samples (Supplementary Figure 7). Thus, for both VFs belonging or not to the T7SS system, we could identify that some genes were absent in some species different from *M. kansasii*, suggesting that these could be part of a specific virulome for each MKC species (Supplementary Figures 6 and 7).

### The MKC antiviral arsenal distribution and diversity

We identified 17 distinct antiviral defense systems among the MKC species (Supplementary Table 7). Wadjet, Restriction-Modification (RM) and Abortive infection (Abi) were the most abundant systems in MKC and were present in all MKC species, with frequencies of respectively 99.6% (662/665), 96.4% (641/665) and 83.2% (558/665) of the genomes. However, while Wadjet and RM were present in most MKC species, Abi was uncommon in *M. attenuatum*, *M. innocens, M. ostraviense* and *M. persicum*. Additionally, in terms of frequency, four other systems (Sirtuin like, Thoeris, Cas, and BREX) were overrepresented in at least one MKC species compared to the others, whereas 10 systems did not present specificity to any species (Supplementary Figure 8). On average, MKC species encode four antiviral systems (3.95), with the average number per species varying from 3.30 to 5.83. The number of antiviral systems per genome varies widely from a minimum of one (as observed in three *M. attenuatum* genomes) to a maximum of eight.

In this study, we also identified 13 CRISPRs regions in the chromosome sequence of *M. kansasii* ATCC 12478; three of those were classified as “confirmed” and ten as “questionable” (Supplementary Table 8). The first confirmed region is located between the genomic coordinates 4428523 and 4428729 and consists of a 26 bp direct repeat (DR) consensus and three spacers. This region seems to be specific to *M. kansasii* as these three spacers were present in all *M. kansasii* genomes and absent in the genomes from other MKC species. The second confirmed CRISPR is the biggest of about 3,3 kbp, starting at 4890197 and ending at 4893369 genomic coordinates, harboring a 26 bp DR and 43 spacers. Spacers of this region were partially or totally shared/absent in MKC species other than *M. kansasii*, possibly representing taxonomic markers. Specifically for *M. kansasii* species, 16 spacers of this CRISPR region were present in all *M. kansasii* genomes, whereas the other 27 were detected in between 3% and 96% of the *M. kansasii* genomes. The third and last confirmed region is placed between genomic coordinates 6379425 and 6380393, comprehends a 24 bp DR and 13 spacers. Spacers of this region were found to be totally present or absent in MKC species other than *M. kansasii*, alike the biggest CRISPR *locus*, and are massively present in *M. kansasii* genomes (≥98%) (Supplementary Figure 9).

### The MKC resistome is species-specific and consists of a wide group of resistance mechanisms

Looking for antimicrobial resistance protein profiles in the chromosome of MKC, we detected 83 resistance protein profiles (Resfams) belonging to 21 resistance mechanisms (Supplementary Table 9, Figure 3). Since the database surveyed encompassed 173 Resfams related to 22 antimicrobial resistance mechanisms, the MKC resistome is extensive, and Nucleotidyltransferase was the unique mechanism without representatives in the MKC species. We observed 57 profiles shared by all MKC species, but some of these profiles have different copy numbers per genome or even average copy numbers per species. Another 10 Resfams profiles are apparently species-related as being absent in at least one MKC species and 16 Resfams were only occasionally observed in some genomes (Supplementary Table 10). Thus, the presence and absence, as well as the variability number of copies, of Resfams within MKC characterize a species-specific resistome.

**Figure 3.**
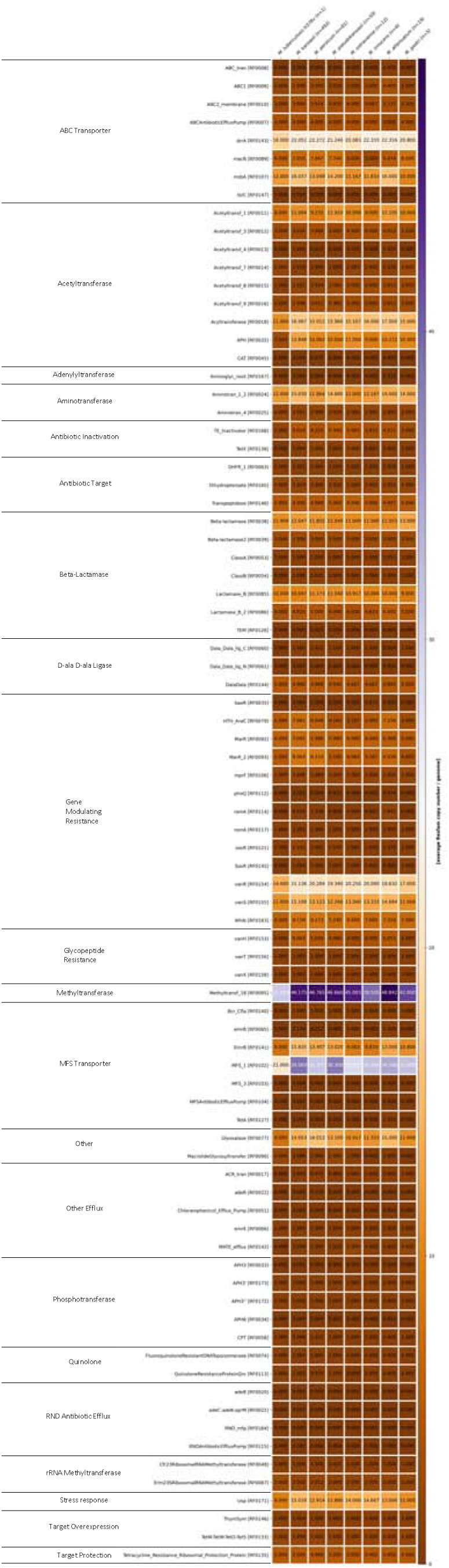
MKC resistome analysis. Heatmaps show the average copy number of resistance protein profiles (Resfams) among each MKC species’ genomes in comparison with M. tuberculosis H37Rv. The graphs show 83 Resfams (second column), classified into 21 resistance mechanisms (first column), found in all MKC species. The average number of copies per genome per species is indicated inside the cells and represented by color degree, according to the legend on the right side of each graph.

By comparing the MKC resistome with the Resfams found in the *M. tuberculosis* H37Rv (n=54), we observe two resistance mechanisms, comprising five Resfams, that are exclusives to MKC (Adenylyltransferase and RND Antibiotic Efflux), whereas the remaining 19 resistance mechanisms were present in both MKC and *M. tuberculosis* H37Rv. The Resfams belonging to those two resistance mechanisms MKC-exclusive were scarcely found in some genomes of the MKC (Supplementary Table 10). Notably, all 54 Resfams found in the *M. tuberculosis* H37Rv are ubiquitous in MKC species, but many of them with distinct copy numbers. In the opposite way, some Resfams not found in *M. tuberculosis* H37Rv are also absent from some MKC species (Figure 3).

We also investigated the resistome encoded by putative MKC plasmids and observed the presence of 27 Resfams related to 11 antimicrobial resistance mechanisms (Supplementary Table 11). Interestingly, a Methyltransferase (RF0095 - Methyltransf_18) was the most commonly observed Resfam in plasmids, the profile also most frequent in chromosomes. However, whereas chromosomes carry several copies of this Methyltransferase, with the average number of copies per species ranging from 39.5 to 48.8, almost all predicted plasmids presented a single copy of this Methyltransferase, except for two *M. kansasii* isolates (KAN-130495 and SRR3666016), having two copies in their plasmid sequences.

### Distribution and emergence of *Mycobacterium kansasii* lineages and sub-lineages

Using Bayesian analysis of population structure (BAPS) to analyze the *M. kansasii* (*sensu stricto*) species population distribution, five distinct genetic groups were defined among the 492 *M. kansasii* genomes. One major group, namely L1, comprises 480 genomes while the other 12 genomes were classified into four small genetic groups (L2, n=4; L3, n=3; L4, n=3; L5, n=2) (Supplementary Table 12). By comparing the genetic groups defined by BAPS with the *M. kansasii* species phylogeny (Figure 4), the groups L1, L2, L3 and L4 characterize four monophyletic lineages comprehending 490 genomes. Two genomes (MK14ES_S84 and KAN-020712) belonging to the L5 BAPS group were divided into distinct branches of the tree, with polyphyletic origin, thus needing further investigation of their evolutionary history.

**Figure 4.**
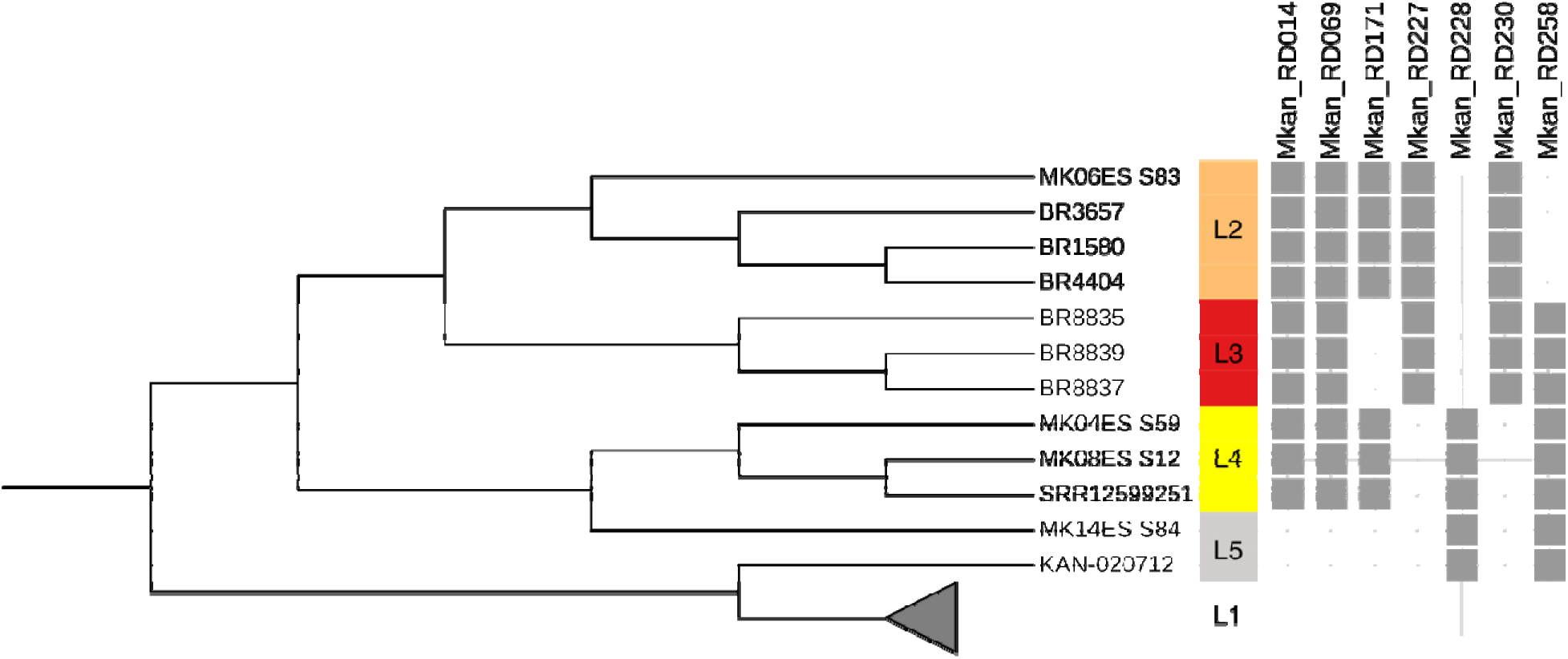
Regions of difference M. kansasii lineage specific. Maximum-likelihood phylogeny tree of 492 M. kansasii genomes, inferred based on nonrecombinant core SNPs, shows the M. kansasii genetic groups defined by BAPS and their specific RDs. Four genetic groups (L1-L4) characterize monophyletic lineages and 2 genomes (MK14ES_S84 and KAN-020712) belonging to L5 genetic group are unclassified due to their polyphyletic origin. Genomes belonging to the lineage L1 (n=480) are collapsed in the tree. Seven regions of difference specific to four (L2-L5) small represented genetic groups are indicated in grey sticks. The RD227 and RD228 shared a genomic region of 4,922bp (genomic coordinates: RD227 from 4766718 to 4771663; RD228 from 4766741 to 4787038), even though RDscan defined as distinct RDs.

Interestingly, from the 12 genomes belonging to L2-L5 BAPS groups, 10 were from Brazil, one was from China (SRR12599251, L4) and one from Belgium (KAN-020712, L3). The main group L1 was further divided into three genetic groups, namely as L1.1, L1.2, and L1.3, with L1.1 as a dominant and comprehending 476 genomes, L1.2 with three genomes from South Korea and L1.3 with one genome from Belgium.

A further analysis of 460 genomes belonging to the main genetic group L1.1 with BAPS defined four L1.1 successful clonal clusters (namely as sub-lineages L1.1.1, n=37; L1.1.2, n=104; L1.1.3, n=186; L1.1.4, n=133). The Bayesian ancestral reconstruction estimated 1815 as most likely date of origin of the root of this main lineage with an evolutionary rate of 4.14e^-02^ (p<1.00e^-04^). An ancestral character reconstruction (Figure 5) estimated the root of the four sub-lineages belonging to L1.1.1, with an unknown geographical origin as there were eight possible countries to root (Belgium, Brazil, China, Czech Republic, Italy, Netherlands, United Kingdom, or United States). The emergence of the other three sub-lineages was estimated as follows: L1.1.2, which include the reference strain ATCC12478, originated from United States in 1929 (marginal probabilities 0.99) and emerging from L1.1.1; L1.1.3 originated from Brazil in 1932 (marginal probabilities 0.82 and 0.99) and also emerging from L1.1.1; and L1.1.4 as the newest sub-lineage, arising in 1966 from Australia (marginal probabilities 0.86 and 0.99) and emerging from L1.1.3.

**Figure 5.**
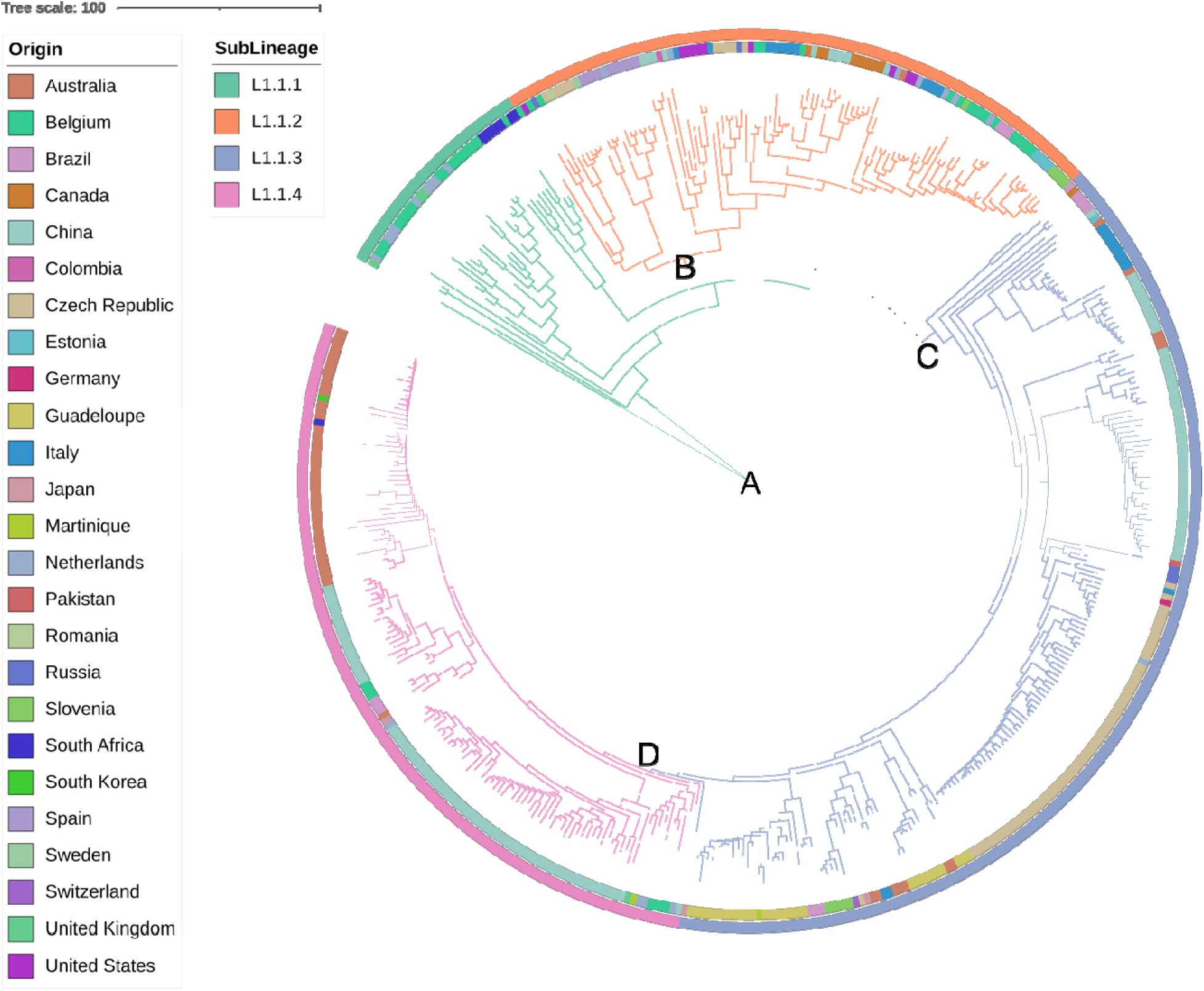
Maximum likelihood dated phylogeny of M. kansasii genomes (n=460) belonging to four sub-lineages. The first inner circle indicates the country of origin of the isolate, wherea the outer circle is the sub-lineage predicted in the Bayesian population analysis. The tree branches are also colored accordingly to sub-lineages, showing the ancestral scenario reconstructed by PastML. The date of the most likely root was 1815, belonging to the older sub-lineage (L1.1.1, n=37) (A), with an unknown geographic region. Three recent sub-lineage emerged in (B) the United States in 1929 (L1.1.2, n=104), (C) Brazil in 1932 (L1.1.3, n=186), and (D) Australia in 1966 (L1.1.4, n=133) (The full tree visualization is also available at iTOL: http://itol.embl.de/external.cgi?tree=157996955338001666969168).

### Fixed mutations in *Mycobacterium kansasii* species could evolve impact on virulence and antimicrobial resistance

We analyzed mutations specific or shared within *M. kansasii* sub-lineages (Supplementary Table 13). Nine mutations were found to be almost exclusive to the older sub-lineage L1.1.1, five mutations shared by the recent sub-lineages L1.1.3 and L1.1.4, and two mutations shared by sub-lineages L1.1.1, L1.1.3 and L1.1.4 were almost totally absent in samples belonging to L1.1.2. Additionally, we observed that 24 positions of the reference strain ATCC12478 presented variants in more than 90% of the *M. kansasii* genomes. Notably, 23 of these 24 mutations did not occur in a *M. kansasii* genome isolated in 1966 (6MK, CP_019885.1), suggesting that the reference positions are not part of the original *M. kansasii* wildtype or that these have been fixed more recently in the majority of the *M. kansasii* population (Supplementary Table 14). Among these 24 variants, four occurred in genes (*CinA*, *embB*, *rpsL* and *rnj*) potentially associated with drug resistance in tuberculosis (TB) and three in genes (*dlaT*, *eccB* and *mycP*) required for virulence activity.

### Regions of difference of *Mycobacterium kansasii* and their possible contribution to the species diversity

In total, 14,621 occurrences of regions of difference (RDs) were observed in the data set of 492 *M. kansasii* isolates (Supplementary Table 15) and were associated to 297 RDs defined by RDscan, with length ranging from 205 to 66,689 bp, and occurring in a range from a single to 442 genomes (Supplementary Figure 10). The three most frequent RDs occurred in over 80% of the genomes and corresponded to two regions with transposases and one region encoding a PE family protein. The largest RD region, corresponding to 66,689 bp, was restricted to five genomes from Guadeloupe (KAN-117, KAN-119, KAN-160, KAN-163, and KAN-165). We also observed that seven RDs occurred only in 12 genomes of four small genetic groups (L2, L3, L4, and L5) (Figure 4). Specifically to RDs restricted to L2-L5, despite RDscan having defined the RD227 and RD228 as distinct RDs, we noticed a shared genomic region of 4,922bp between these two RDs (genomic coordinates: RD227 from 4766718 to 4771663; RD228 from 4766741 to 4787038), causing the loss of four coding sequences (MKAN_RS20805: “RtcB family protein”; MKAN_RS29915: “AAA family ATPase”; MKAN_RS20830: “XRE family transcriptional regulator”; and MKAN_RS20835: “type II toxin-antitoxin system RelE/ParE family toxin”), one tRNA (MKAN_RS20815) and one pseudogene (MKAN_RS20840). Therefore, the shared region of RD227 and RD228 likely characterize the distinction between the large lineage L1 and the other genetic groups L2-L5. There was a distinction in the RDs average per genome for the whole dataset (29.78, SDL=L20.29, Median = 25) when compared with the four defined for sub-lineages (L1.1.1 = 21.35, SD = 13.09, Median = 18; L1.1.2 = 21.76, SD = 16.62, Median = 17; L1.1.3 = 25.30, SD = 20.28, Median = 16; L1.1.4 = 40.89, SD = 18.05, Median = 43) (Supplementary Figure 11).

### Analysis of *Mycobacterium kansasii* from the Czech Republic indicates a higher transmission rate for the sub-lineage L1.1.3

We further analyzed a collection of 68 *M. kansasii* isolates from the Czech Republic collected in 1996 and 1997, in order to understand if some of them belong to the same outbreak and their possible transmission clusters (Supplementary Table 16). These isolates were either clinical (n=62) or environmental (n=6), the latter derived from artificial water sources in four distinct coal mines. Among the clinical samples, 15 were collected from employees of three of these mines. Additionally, from the 62 clinical isolates, 57 patients were considered with NTM disease, four colonizers, and one with unknown clinical phenotype. The genotyping showed that isolates belonged either to sub-lineages L1.1.2 (n=10) or L1.1.3 (n=58), suggesting they not belong to the same outbreak. The SNP distance between isolates ranged from 0 to 135, with a mean SNP distance between all Czech isolates was 41.55 (SD=26.78). Considering only isolates belonging to L1.1.2 sub-lineage the mean was 52.93 (SD=24.47), and 28.70 (SD=16.11) for isolates belonging to the L1.1.3 sub-lineage. The smaller distance for isolates from distinct lineages was 56, two pars of isolates had zero SNP distance (KAN-961342 and KAN-961343, belonging to L1.1.2; and KAN-961384 and KAN-961392, belonging to L1.1.3). Considering the pairwise distance between each Czech isolate with those whose belong to the same sub-lineage, the mean for L1.1.2 Czech isolates ranging from 41.22 to 71.66 whereas for L1.1.2 Czech isolates were between 15.05 and 68.03. Interesting, the sample with lowest mean distance (15.05) is an environmental sample (KAN-970587) isolated from a shower-bath in a coal mine. By applying the SNP distance thresholds of 10, 20 and 30, the number of possible transmission clusters identified were, respectively, five, three and four. Notably, regardless of the SNP threshold applied, these clusters include clinical and environmental isolates and colonizers (Figure 6). Upon considering the geographic distribution of the lineages, we observed some genotypes were from isolates derived from patients of distant geographic origin.

**Figure 6.**
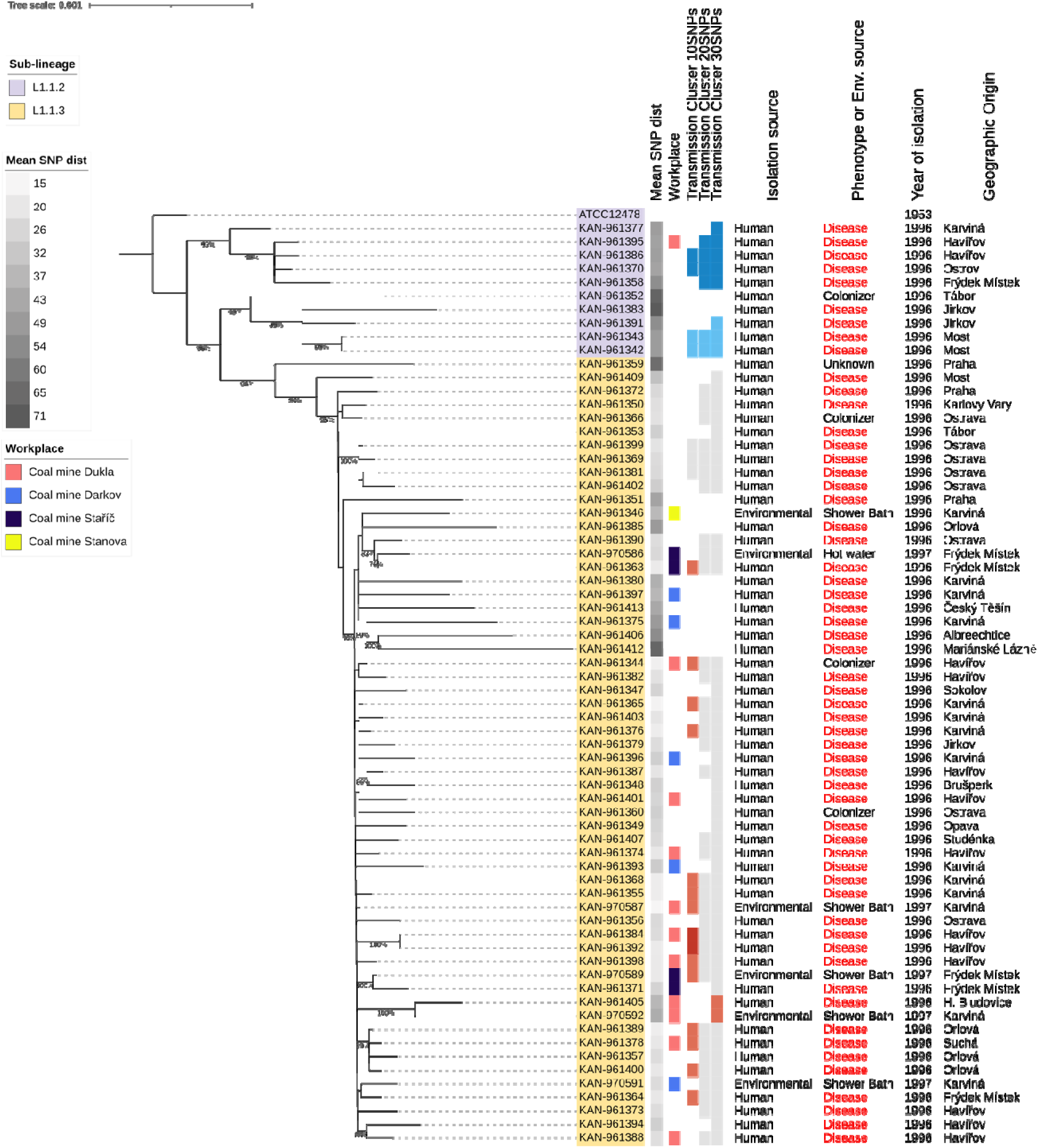
Maximum likelihood phylogeny tree of M. kansasii strains isolated in Czech Republic. The tree inferred based on nonrecombinant core SNPs is rooted in the reference strain ATCC12478. The number of nucleotide substitutions per site is indicated by the scale bar and bootstrap values are indicated in branches with at least 75% of support. Sample sub-lineage is indicated by colors at node names. Beside the tree, from left to right, the following characteristics are described: the mean SNP distance between a sample and their sub-lineage group; the workplace (where available); bars with colors indicating transmission clusters inferred for three distinct SNPs cut-offs; source of the isolation as either human or environmental; clinical phenotype (human isolates) or detailed isolation source (environmental isolates); year of isolation; and geographical origin of the isolate (environmental isolates) or patient (human isolates).

## Discussion

The findings of this study highlight several genomic characteristics of the non-tuberculous mycobacterium *M. kansasii* complex, demonstrating that the seven known MKC species harbor unique and shared genomic signatures, more specifically in their pangenome, resistome, virulome, mobilome and defense systems. We expanded the publicly available MKC data by providing genome sequencing of 342 isolates and, in line with previous reports^1,2,8^, our data support the known MKC population distribution into seven distinct species. Additionally, we have provided new MKC WGS data for isolates from 28 countries, including 10 countries with no MKC available data until now (Bangladesh, Colombia, Estonia, Guadeloupe, Martinique, Pakistan, Romania, Russia, Slovenia, and Sweden). After realizing our experiments, Zhang et al.^15^ and Rajendran et al.^16^ published data including MKC genomes from respectively 237 and 12 isolates from China and India.

Our results further illustrate that the MKC accessory genome is diverse and the MKC pangenome is still open and expansive with species-specific accessory gene content. Additionally, this illustrates that mobile genetic elements contribute to the acquisition of new genes by MKC species, as documented by the high frequency of strains with plasmid and prophage signatures. We show that these pangenome characteristics remain observable regardless of the similarity cut-off applied, although 80% percent is likely a good cut-off for the MKC pangenome definition.

The variability of virulence factors within MKC species is supported by earlier data, comparing *M. kansasii* against other species^6,17,18^ suggesting an MKC species-specific virulome. To address this, we defined 287 *M. kansasii* genes orthologs to *M. tuberculosis* H37Rv potential virulence factors and screened for their presence in the genomes collection. We observed unique and shared virulome characteristics among the MKC species with 123 VFs ubiquitous to MKC and 164 genes partially or totally absent from some MKC species or strains. In concordance with their frequent isolation from diseased humans, *M. kansasii* and *M. persicum* were the species with the highest number of virulence factors. Our virulome analysis also encompasses genes belonging to the type VII secretion systems (T7SS), which consists of five paralogous loci (ESX-1 to ESX-5) in *Mycobacterium tuberculosis* and is known to play a crucial role in mycobacterial survival in the host, phagosome escape and immune response evasion^19^. Looking for orthologs of the *M. tuberculosis* H37Rv T7SS genes, the majority of *M. kansasii* isolates presented all orthologs as previously observed^9^, while only two orthologs (*espG1* and *espI*) being absent in *M. persicum*. For the remaining species, the number of H37Rv T7SS orthologs not identified varies between six and 20. This again may explain the lower association of human disease and species other than *M. kansasii* and *M. persicum*.

Encoded outside the T7SS gene cluster, the *espACD* operon is recognized as essential for ESX-1 secretion^20^. In disagreement with a previous report^17^, we observed that the presence of the *espACD* operon is not exclusive for *M. kansasii* and was also present in all *M. pseudokansasii* and *M. attenuatum* genomes. Considering that our data include several *M. persicum* isolates from clinical cases, this indicates that the *espACD* operon is not a unique feature for MKC species pathogenicity. Notably, one *M. kansasii* isolate (KAN-43) lacked the *espACD* operon, with a deletion of 6,874bp that was also confirmed using RDscan analysis (Mkan_RD151). Still, with respect to the species or strains virulome characteristics, even though each MKC species generally keeps a specific set of VFs, some strains are deviating from their species virulome gene set, by losing some VFs. This poses that distinct strains of MKC species can present a broad virulence spectrum, as already observed for *M. kansasii*^6^. Interestingly, KAN-43 is one of the two strains that belonged to phage type 2 and if the absence of the *espACD* operon has influence on the straińs virulence is currently under investigation.

We have proposed a holistic approach to address the MKC resistome by looking for resistance protein profiles (Resfams) across the analyzed genomes. Similar to the MKC virulome, the MKC resistome is species-specific, and a high number of Resfams (n=83) within MKC species were identified, belonging to 21 of the 22 resistance mechanisms described. This suggests a large potential for resistance against xenobiotics and seems compatible with an organism lifestyle capable of growing in different environmental conditions. The MKC resistome is also larger than that of *M. tuberculosis* H37Rv and in general presents more copies of resistance profiles ubiquitous to both MKC species and H37Rv. The variability of Resfams copy numbers corroborates that distinct MKC species can have different antimicrobial phenotypes, specially related to intrinsic antimicrobial resistance levels. Our observation of the presence of 27 Resfams within MKC plasmids strengthens the hypothesis of expansion of the MKC resistome by lateral transfer events. Treatment for MKC infections resembles the TB therapy scheme and is a challenge due to the long period and demand for multiple drugs^21^. Hence, we believe that this MKC resistome panorama and comparison to that of *M. tuberculosis* H37Rv can support further analysis of the MKC species resistance to antimicrobials, as well as possible changes to the current standard of care therapy for MKC infections.

We have identified putative plasmid-derived contigs within 36.7% of the MKC genomes, slightly more than 29.3% previously observed^7^. The larger frequency observed is probably due to our combined approach of three *in silico* strategies, as well as to the use of larger databases, not restricting the search to a few genetic markers and avoiding a selection bias, as such, expanding the knowledge on plasmids diversity within MKC genomes. All genomes with plasmids had at least one replicative, conjugative or mobilizable plasmid marker identified. As expected, homologs of the Type IV and Type VII secretion systems are common among MKC plasmids and besides their potential contribution to virulence, contribution to antimicrobial resistance was likely because of the presence of Resfams profiles.

Except for *M. gastri*, we found that prophages are common to MKC genomes. As the few known *M. gastri* isolates with known origin come from humans, this species may be host-restricted, thus indicating a likely reason for the absence of prophages, due to limited exposure to phages and evolutionary adaptation. Notably, despite most MKC prophages and other know mycobacteriophages are related as evidenced by their shared genes connections, all predicted MKC prophage clusters were distinct from clusters of known phage representative sequences. Similar to the *M. abscessus* complex^22^, our results suggest that phages can be exchanged by different MKC species, as many defined prophage clusters were shared by distinct MKC species. However, this does not exclude species phage affinity because some prophage clusters were species-specific. To what extent these prophages contribute to biological activity and are related to active phages needs further investigation.

As an extension to our analysis of presence of prophages, we identified the MKC arsenal of defense systems mechanisms and observed that most MKC genomes contain three distinct defense mechanisms. One is of the Wadjet type, conferring protection against plasmid transformation^23^; the second is the RM type, that acts by recognize and target sequences of bacteriophages^24^; and finally, the Abi type, an abortive system that, upon infection, leads towards cell death or metabolic arrest^25^. Besides these three, even though less frequent, another 14 defense systems were identified and, showing the potential of different MKC species to acquire distinct anti-phage defense mechanisms. However, given the observed variability and diversity of defense mechanisms and the limited number of isolated from some of the MKC species available, the acquisition and maintenance of such systems in MKC genomes must be further investigated. In addition, we identified three different CRISPR regions along the reference genome of the strain *M. kansasii* ATCC12478, including the identification of spacers with variable presence along strains and species of the complex. This points to possible differences in specificities of defense against phages and could lead to the development of a typing system such as spoligotyping used for the *M. tuberculosis* complex ^26^. Interestingly, CAS defense systems were inferred in only 5% of *M. kansasii* genomes and not detected in the ATCC12478 strain. So, these CRISPR regions are isolated from a *cas* loci and do not constitute a CRISPR-CAS system, as observed in other bacterial genomes^27^.

Besides confirming the seven main species within the MKC in the largest dataset analyzed so far, our findings demonstrate that the *M. kansasii* species can be subdivided into four monophyletic lineages, which was similar as previously observed^13,18^. One of these, the major lineage (L1), is globally distributed and comprehending four successful clonal groups (sub-lineages L1.1.1, L1.1.2, L1.1.3 and L1.1.4). We observed that Brazil is as an important source of *M. kansasii* diversity, as 10 of 12 genomes from the scarce represented genetic groups L2-L5 had been isolated in the country. This genetic diversity of *M. kansasii* isolates from Brazil has been previously suggested^13^, but it is remarkable that from almost 500 isolates this group of few diverse genomes is mostly restricted to Brazil. This urges the need to investigate more isolates from this country for better understanding of the L1 local and global phylogeny. We evidenced that three (L1.1.2, L1.1.3 and L1.1.4) of the four most represented and successful sub-lineages emerged during the 20^th^ century, with the newest sub-lineage since the 1960s. The highest frequency of deletions was observed in the most recent sub-lineage L1.1.4. Additionally, we observed that *M. kansasii* genetic groups L2-L5 are characterized by a region of difference (RD) of 4,922 bp, resulting in loss of one tRNA, one pseudogene and four coding sequences that include a RtcB protein, an ATPase, a XRE transcriptional regulator and a type II toxin-antitoxin RelE/ParE.

Regarding genetic differences among *M. kansasii* we observed SNPs found in all or almost all genomes analyzed and some of these mutations were observed in genes with potential impact on virulence (*dlaT*, *eccB*, and *mycP*) and antimicrobial resistance (*CinA*, *embB, rpsL,* and *rnj*) ^28–33^.

Our collection of strains from Czech Republic provided data to infer *M. kansasii* transmission cluster analysis. Importantly, even for a restrictive clustering threshold (distance of 10 SNPs), our study indicates the presence of *M. kansasii* colonizers, clinical and environmental isolates belonging to the same cluster, suggesting potential to both human-to-human and environmental-to-human transmission.

## Methods

### Data, sample collection, and whole-genome sequencing

The present study includes a total of 342 isolates from 25 different countries, encompassing five continents. Briefly, 298 isolates were obtained from human clinical samples, 45 from environmental niche, 41 with unknown origin, and four from animals, collected between 1996 to 2016. These isolates are derived from culture colonies presenting MKC-like phenotypic characteristics and were subjected to whole genome sequencing. NGS libraries were constructed from genomic DNA using the Nextera protocol^34^ and the Illumina NextSeq 500 platform with 2 x 151 bp runs (Illumina, San Diego, CA, USA). We additionally sequenced with PacBio long-read sequencing on an RSII instrument (Pacific Biosciences, Menlo Park, CA, USA) three MKC isolates (KAN-C06235, KAN-130495 and KAN-960446) to generate complete or near fully complete closed genome sequences, as these strains had enough DNA of sufficient quality to PacBio sequencing, in order to decipher their whole plasmid content. In addition to our data collection, we retrieved available MKC sequence read data sets and assembled genomes from NCBI nucleotide databases as of May 2021, excluding next-generation sequencing (NGS) data from non-Illumina platforms. Public sequence read data was downloaded from SRA database and converted into FASTQ files using the NCBI SRAtoolkit (v2.10.8, https://ncbi.github.io/sra-tools/).

### Genome assembly and annotation

Illumina sequencing reads were trimmed and filtered using Trimmomatic (v0.36)^35^ and genomes were assembled with SPAdes (v3.14.1)^36^ applying the careful mode, auto coverage cutoff, mismatch corrections, and automatic selection of the k-mers size based on read length. Contigs of less than 500 bp and coverage lower than 2x were discarded. PacBio genomes were assembled with CANU (release 10117)^37^ applying the following parameters: *genomeSize=6.57mb* for all samples, *minReadLength=2000* for sample KAN-960446, *minReadLength=3000* for sample KAN-C06235, and *minReadLength=4000* for sample KAN-130495. All assemblies were assessed for quality using the CheckM lineage workflow (v1.1.2)^38^ and the average nucleotide identity (ANI) was calculated for all genome pairs using fastANI (v1.31)^39^ with the option ‘many-to-many’ and default parameters. We excluded assemblies with deviating genome number of contigs >L1,000, less than 95% pairwise ANI with any of the MKC reference genomes (*M. kansasii* ATCC12478, CP006835; *M. persicum* AFPC-000227, MVIF01; *M. pseudokansasii* MK142, UPHU01; *M.ostraviense* 241/15, NKRE01; *M. innocens* MK13, UPHQ01; *M. attenuatum* MK41, UPHT01; *M. gastri* DSM43505, LQOX01), and with more than 10% of contamination as estimated by CheckM. The resulting data set contained 665 genomes (Supplementary Table 2), in which 342 were generated in this study and 323 retrieved from public databases. Genomes were annotated with Prokka (v1.14.6)^40^.

### Pangenome inference and core genome definition

The MKC pangenome was calculated and visualized using Roary (v3.13.0)^41^. Since the MKC harbor seven species, we generated three pangenome runs applying different BLASTp identity cutoffs (parameter *-i* setted to 90%, 80%, and 70%), to verify the impact in the core genome estimation. The remaining Roary parameters were the same for all runs, as follows: *-s* (no paralog splitting), *-cd 99* (a gene must be in 99% of isolates to be core) and *-e --mafft -n* (core gene alignment generated with MAFFT (v7.487)^42^. Pangenome matrixes were visualized with Phandango^43^.

### Phylogenomic analysis

For the MKC phylogenetic analyses, we generated a concatenated alignment of the core 1358 shared genes, that was created with Roary applying the following parameters: “*-i 95 -s -cd 100 -e --mafft -n*”. Thus, a starting tree was built using RAxML-NG (v1.0.1)^44^ applying a GTRLmodel,LGamma correction and 100 bootstrap replicates. The same genome alignment was fed into fastGEAR (v1.0.1)^45^ to detect recombinant regions. These regions were then parsed into a text file storing recombinant genomic regions as coordinates (as a BED file format - Browser Extensible Data) to generate a recombination masked alignment using maskrc-svg (first release, https://github.com/kwongj/maskrc-svg) and the starting tree. Hence, the masked alignment was used to build a recombination-free maximum-likelihood (ML) phylogeny in RAxML-NG with the same approach as above plus ascertainment bias correction, using the Lewis’ method (parameter +ASC_LEWIS). The final tree was employed and annotated in iTOL (v5)^46^. The core gene alignment length encompassed 334,144 sites, amounting to 222,205,760 characters for the whole data set. Across all isolates, 85,633,543 positions (38.53% of all core gene alignment sites) were masked for recombination. For the final recombination-free ML phylogeny analysis, all invariant sites were removed with snp-sites (v2.5.1)^47^ to obtain the final alignment length of 48,820 variants.

The *M. kansasii* species (former subtype I), the *M. kansasii* lineage L1.1 (460 genomes with available DNA sequencing reads, Supplementary Table 12), and the *M. kansasii* Czech isolates phylogenies were inferred based on genome alignments generated with Snippy (v4.6.0, https://github.com/tseemann/snippy). Alignments were analyzed with Gubbins (v2.4.1)^48^ to detect and mask recombinant regions. After, invariant sites were removed with snp-sites and the resultant alignments were then used to detected phylogenetic lineages or sub lineages and to generate a ML phylogenies with RAxML-NG, applying a GTR model,LGamma correction, Lewis’ method ascertainment bias correction and 100 bootstrap replicates. Lineages and sub lineages were detected by applying a hierarchical Bayesian Analysis of Population Structure (hierBAPS) implemented in R (v1.0.1)^49^ with a maximum depth of two and 50 as a maximum population number. Additionally, the Gubbins output analysis of the *M. kansasii* lineage L1.1 was used as input to infer a dating phylogenetic tree with a Bayesian approach, by using BactDating (v1.1)^50^ with 10^6^ iterations. The *M. kansasii* lineage L1.1 dated tree fed PastML to the ACR inference for lineage and geographical origin, applying the maximum likelihood prediction method MPPA (marginal posterior probabilities approximation) and evolutionary model F81. The pairwise SNP distance for *M. kansasii* Czech isolates was evaluated using snp-dists (https://github.com/tseemann/snp-dists). *M. kansasii* Czech transmission clusters were defined using the library “cluster” with R statistical programming language and applying thresholds of 10, 20 and 30 SNPs.

### Detection of mobile genetic elements

The identification of contigs with a possible plasmid origin was performed by using Platon (v1.6)^51^, viralVerify (v1.1, https://github.com/ablab/viralVerify), and searches with BLASTn^52,53^. For the BLASTn approach, we compared MKC contigs against a customized database with 36,425 prokaryotic reference plasmids available on the RefSeq database (release 207) together with complete plasmid sequences of three MKC genomes (strains MK142, KAN-960446, and KAN-C06235). We applied the following parameters: e-value 1e^-05^, 80% of identity, and word size = 11. We considered a contig as originated from plasmid as follows: (i) if a contig was classified as plasmid by both Platon and viralVerify; or (ii) if a contig was classified as plasmid only by one software (Platon or viralVerify) and had a BLASTn hit with a reference plasmid; or (iii) if Platon classified the contig as plasmid due to a hit with a mobilization protein profile and viralVerify resulted in “uncertain”, regardless without a BLASTn hit, as Platon is based on high-quality Hidden-Markov Model (HMM) profiles of relaxase protein families available on the MOBscan database^54^. To avoid false positives, we discarded contigs with a sequence size less than 5 kb and BLASTn hits with less than 50% of coverage. Besides the Platon prediction of hits with reference replication, mobilization, and conjugation proteins in the predicted plasmids, we additionally sought for the occurrence of marker proteins of the reference plasmid pMK12478 (NC_022654). This survey was performed by extracting from viralVerify results hits with type VII (T7SS) and type IV (T4SS) secretion systems proteins, based on HMM profiles of protein families (Pfam) listed in Supplementary Table 17.

Prophages were detected with VirSorter2 (v2.2.3)^55^ and CheckV (v0.8.1, database v1.0)^56^. First, genome sequences were surveyed with VirSorter2 applying the parameters *--include-groups dsDNAphage*, *--min-length 5000*, and *--min-score 0.75*. After, we checked the VirSorter2 resultant fasta files (*final-viral-combined.fa*) with the *end_to_end* workflow of CheckV. Finally, the *proviruses* and *viruses* fasta files resultant from the CheckV analysis were submitted over again to VirSorter2, with the same parameters of the first step analysis. Sequences of the inferred prophages were annotated with DRAM-v (v1.2.4)^57^. Thus, to identify possible prophages species specific as well the similarity between MKC prophages and known Mycobacteriophages, a bipartite network and clusters of the prophages and their gene content (putative encoded proteins) were generated with AccNET (v1.2)^58^, employing the parameters *--threshold 1.12* and *--kp* ′*-s 1.8 -e 1e-5 -c 0.6*′. For the clustering and network analysis performed by AccNET we also included 161 representatives Mycobacteriophages retrieved from PhagesDB^59^ (http://phagesdb.org, accessed in August 2021 - Supplementary Table 18), that encompass all mycobacteriophage clusters available on the PhagesDB. The cut point 0.90 was applied for clustering heights, as this quantile was able to define almost exactly the clusters available on the PhagesDB, and the resultant network was employed in Cytoscape (v3.7.2)^60^.

### Resistome and virulome analysis

The survey of genes evolved with possible antibiotic resistance activity was based on Hidden Markov Models profiles of the ResFams database^61^, using HMMER (v3.3.2)^62^ and applying an e-value 1e^-09^.

Genes with putative contribution to virulence activity were surveyed in MKC genomes with Abricate (v1.01, https://github.com/tseemann/abricate), based on a custom database with 287 *M. kansasii* genes with recognized orthologs in *M. tuberculosis* H37Rv that encode for potential virulence factors found by Mussi et al.^6^. We applied a cutoff of 85% identity and coverage.

### Antiviral defense systems identification

We define the presence of Antiviral defense systems based on PADLOC (v1.0.3, database v1.2.0)^63^ and DefenseFinder (first release)^64,65^. Firstly, systems were assigned within MKC genomes applying the PADLOC results and the defense systems BREX, Dnd, Paris, and RM were additionally inferred by DefenseFinder. Additionally, we sought for CRISPRs regions on the *M. kansasii* ATCC 12478 genome with CRISPRfinder online tool (https://crispr.i2bc.paris-saclay.fr/Server/, database version 2017-05-09)^66^, and regions identified as “confirmed CRISPRs” were blasted against all MKC genomes analyzed in the work.

### Structural variants within *M. kansasii* species

We investigated regions of difference (RDs) within genomes of the *M. kansasii* species by adapting RDscan (first release)^67^ to map DNA sequencing read data of the *M. kansasii* isolates against the *M. kansasii* ATCC12478 chromosome (NCBI version NC_022663.1). Once RDscan is based on a read mapping approach, we generated fake reads for samples obtained from NCBI at genome level, without publicly available DNA sequencing reads. The following RDscan analysis parameters were applied: *threshold: 0.05, DHFFC: 0.1, minSVLEN: 200,* and *maxSVLEN: 100000*. Fake reads were generated with readSimulator (first release, https://github.com/wanyuac/readSimulator), applying Wgsim (first release)^68^ to generate synthetic paired-end short reads under the parameters: *--iterations 10 --readlen 200 -- depth 100 --opts ’-e 0 -r 0 -R 0 -X 0 -h -S 5’*.

### Phage typing

Susceptibility to 14 phage types of the MKC strains at the RIVM (National Institute for Public Health and the Environment, Bilthoven, The Netherlands) was determined as described by Engel and Berwald^69^.

## Supplementary material

Supplementary tables 1 to 19 providing details of used data and results of this study are available in the Supplementary Data 1. Supplementary Figures 1 to 5 and 8 to 11 are provided in the Supplementary Data 2. The Supplementary Figures 6 and 7 are, respectively, available in the Supplementary Data 3 and Supplementary Data 4 files.

## Author contributions

E.M., P.S. and C.J.M. conceived the study, curated the data, visualized the results and wrote the initial draft. E.M., S.V., L.G., M.C., L.R., R.J., S.C., P.R., C.U., T.L., H.N. and P.S. performed the formal analysis. J.R., L.C., T.G., M.D., P.R., P.C., C.L., L.S., S.V., J.K., N.R., K.L., I.M., V.Z., N.M., M.Z., V.J., J.W. and M.S. provided sample data. C.J.M. and P.S. acquired funding. C.J.M., P.S., D.S., S.N. and B.C.J. provided resources for this study. All authors critically reviewed and modified the paper.

## Acknowledgments

We are grateful to Plataforma de Bioinformática–RPT04A (Rede de Plataformas Tecnológicas FIOCRUZ) by provide support and infrastructure to bioinformatic analyses.

