## Supplemental Data 2 for "Phylogenomic and genomic analysis reveals unique and shared genetic signatures of *Mycobacterium kansasii* complex species"

**Supplementary** Figure 1

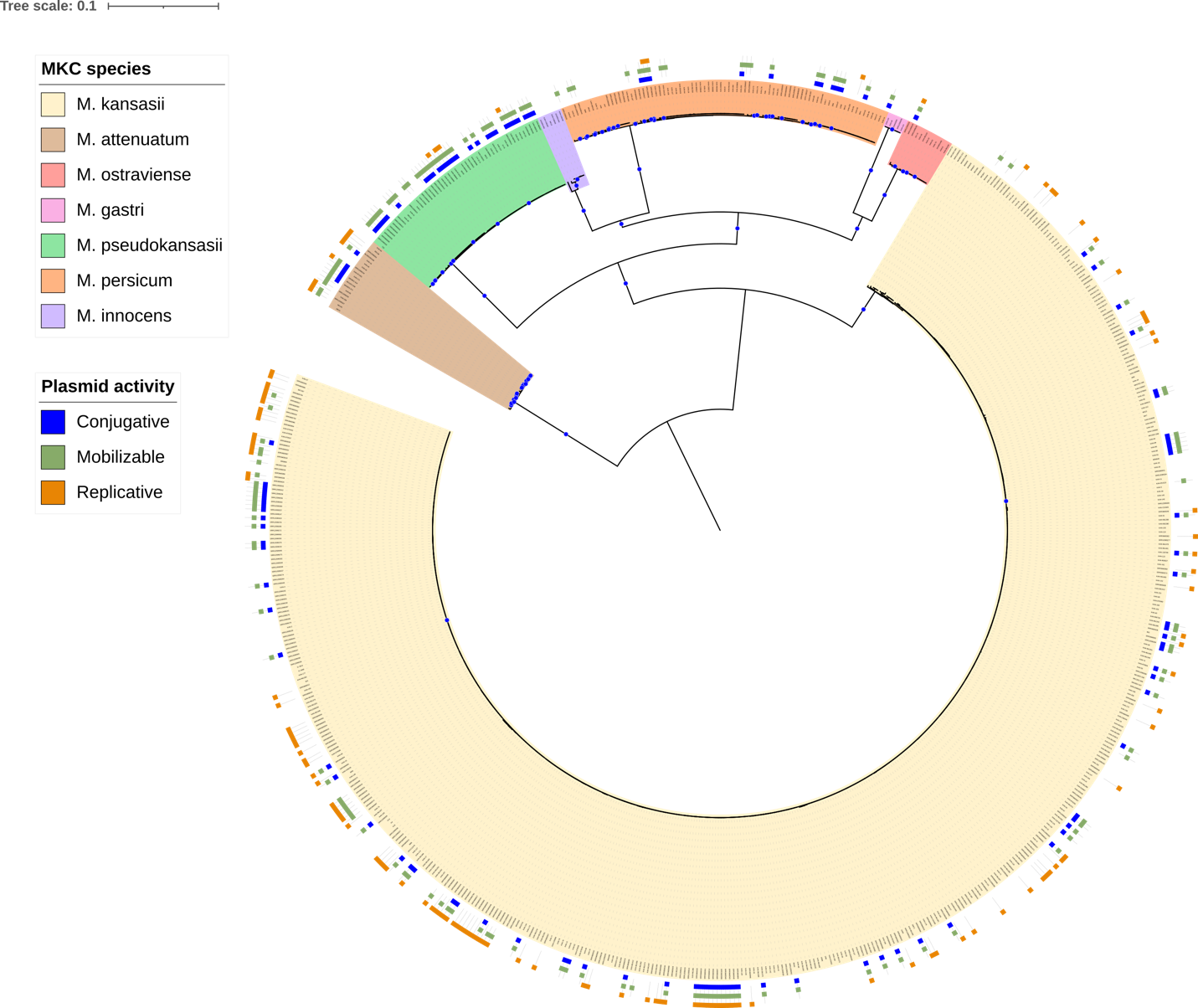

**Supplementary Figure 1**. **Plasmids distribution among MKC**. Midpoint rooted maximum likelihood phylogenetic tree showing the prevalence and distribution of plasmids among 665 *M. kansasii* complex genomes. Blue dots indicate nodes with bootstrap support above 75% with shading of branches and nodes indicating the seven MKC species. The three outer rings denote the plasmids machinery activity inferred, as either conjugative, mobilizable or replicative.

**Supplementary** Figure 2

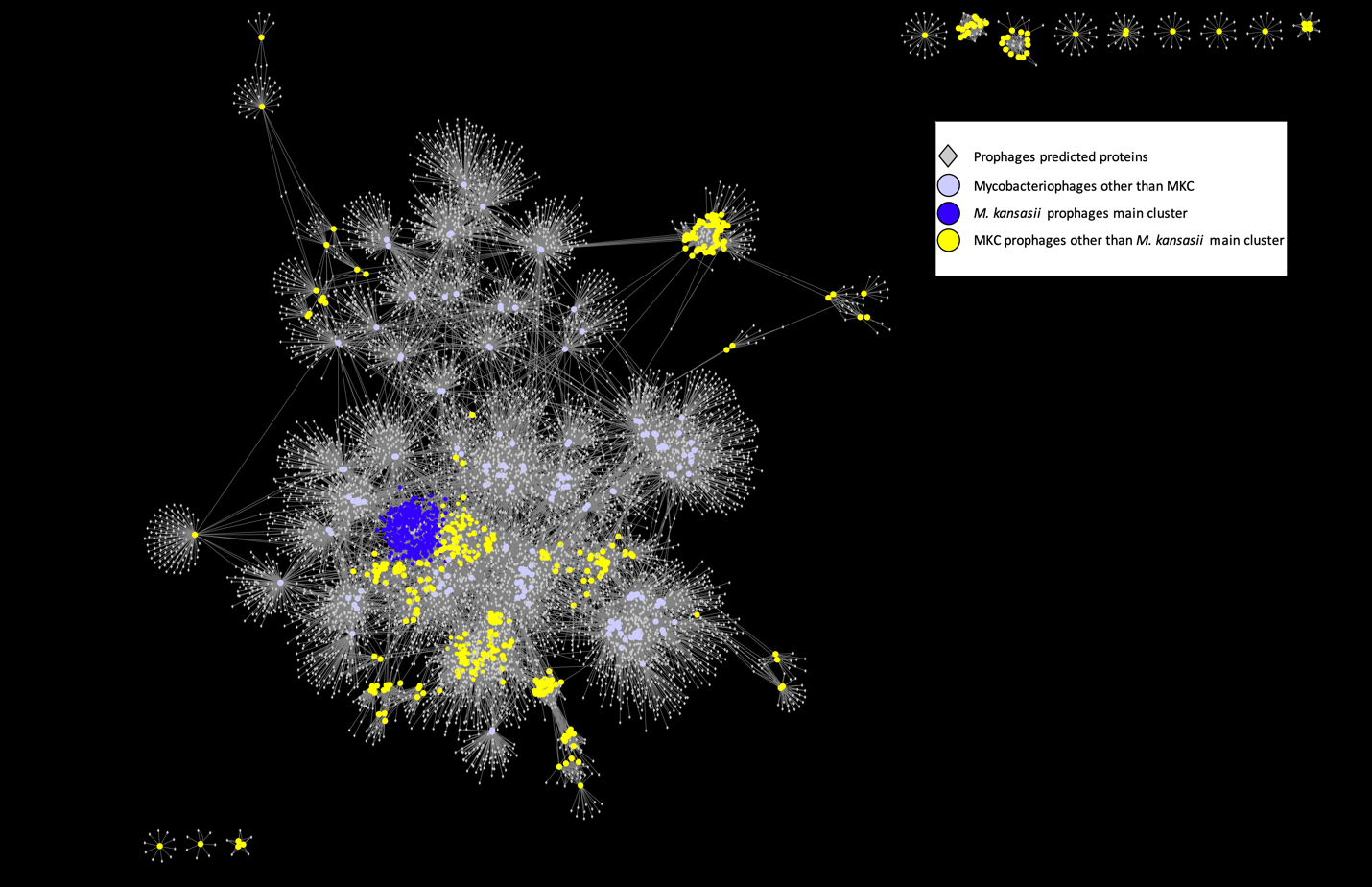

**Supplementary Figure 2**. **MKC prophages network**. The bipartite network of the prophages and their gene content. The predicted genes of the prophages are show as grey diamonds, whereas prophages genomes as colored circles. (A) Genomes circles are colored by MKC species or Mycobacteriophages genomes obtained from hosts other than MKC species, as defined in the right upper legend. Genomes are connected to their predicted genes by edges. (B) Genomes belonging to the main *M. kansasii* cluster, with 335 similar prophages, are show as dark blue circles. MKC prophages genomes belonging to clusters other than the main *M. kansasii* prophage cluster and Mycobacteriophages genomes are, respectively, represented by yellow circles and light blue circles.

**Supplementary** Figure 3

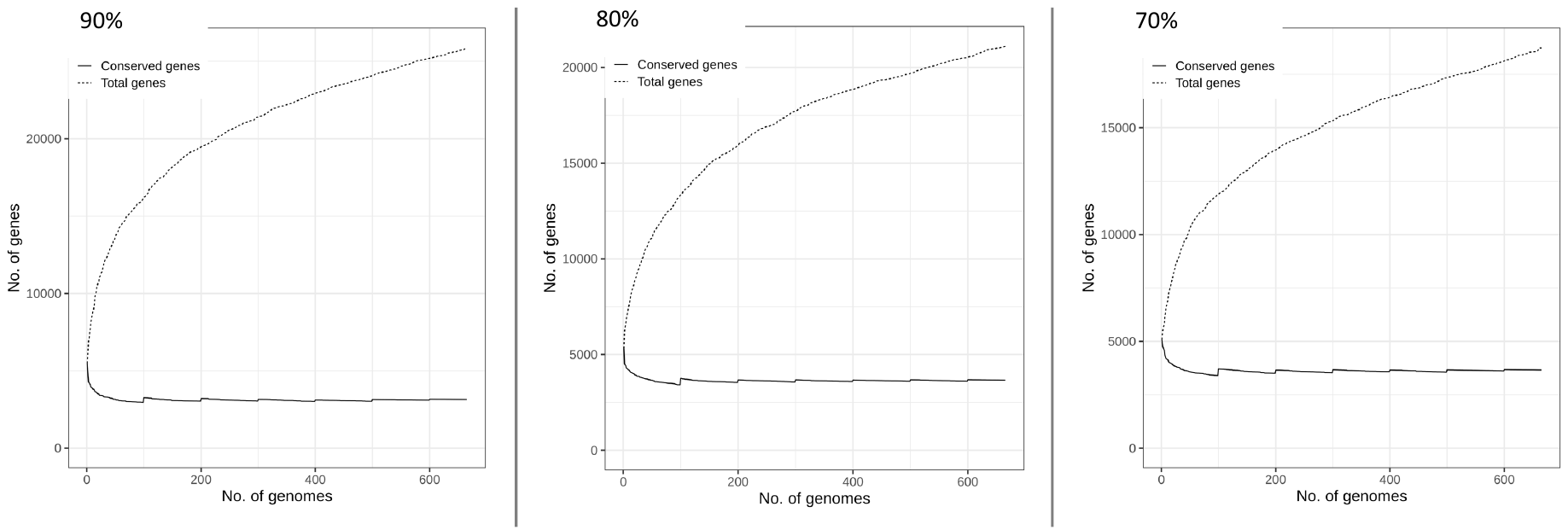

**Supplementary Figure 3**. **MKC pangenome curve** for three runs applying different similarity cut-offs (90%, 80%, 70%). Each graph axis represents the number of genes (vertical) *versus* the number of genomes (horizontal) and shows two curves: the total number of genes (dot line) and conserved genes (continuous line) in the pangenome. Regardless of the similarity cut-off, the number of total genes does not achieve a plateau, thus characterizing an opened pangenome.

**Supplementary Figure 4**

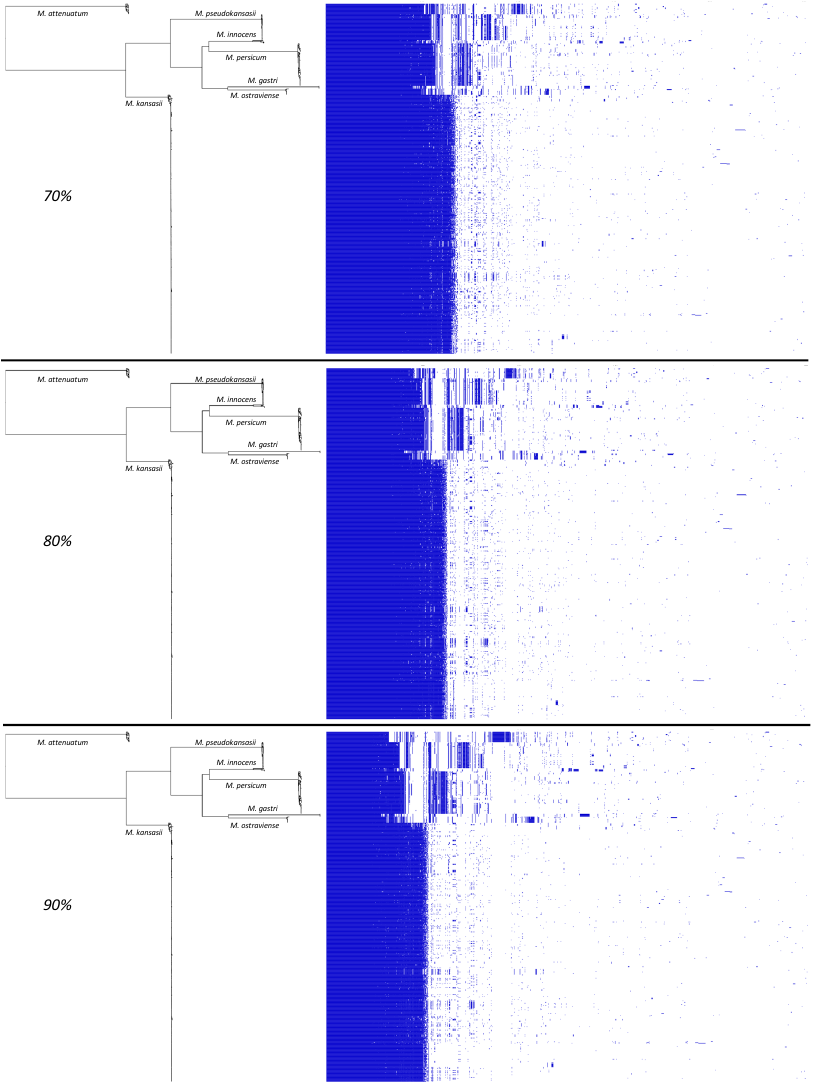

**Supplementary Figure 4**. **MKC pangenome matrix**. The *M. kansasii* complex accessory gene content is structured by species regardless the similarity cut-off applied (70%, 80% or 90%, from top to bottom). The pangenome matrixes shows the presence (blue) or absence (blank) of homologues genes among MKC species genomes are plotted (right), and the phylogenetic trees (left) corresponds to the tree in Figure 1.

**Supplementary Figure 5**

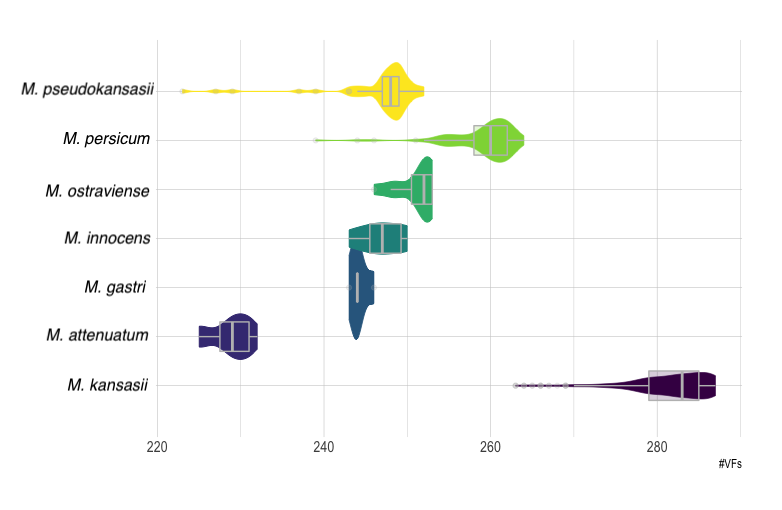

**Supplementary Figure 5**. **MKC virulome characteristics**. Violin plot showing the number of virulence factors (x axis) for each MKC species (y axis). Density peaks of virulence factors inferred are shown in the graph for each MKC species. Additionally, internal box plots shows summary statistics (median and interquartile). The thin gray line represents the rest of the distribution, whereas points are outliers.

**Supplementary Figure 6** (available in PDF). **Presence or absence of the Type VII secretion system (T7SS) genes among *M. kansasii* complex genomes.** Midpoint rooted maximum likelihood phylogenetic tree of 665 *M. kansasii* complex genomes and the presence or absence of the Type VII secretion system (T7SS) genes. Node names are colored by MKC species, as indicated in the left upper legend. Genes are organized in the five paralogous ESX loci (ESX-1 to ESX-5), with the reference *M. kansasii* ATCC12478 locus name in brackets. Additionally, the respective *M. tuberculosis* H37Rv orthologous gene name and locus are given in parenthesis. Circles represents the presence (black filled) or absence (unfilled).

**Supplementary Figure 7** (available in PDF). **Presence or absence of 119 virulence factors among *M. kansasii* complex genomes**. Midpoint rooted maximum likelihood phylogenetic tree of 665 M. kansasii complex genomes and the presence or absence of 119 virulence factors (VFs). Node names are colored by MKC species, as indicated in the left upper legend. The presence (filled shapes) or absence (unfilled shapes) of VFs for each genome is indicated. Functional category of VFs is indicated by shape forms and colors. The reference *M. kansasii* ATCC12478 locus name is given in brackets, whereas the respective *M. tuberculosis* H37Rv orthologous gene name and locus are given in parenthesis.

**Supplementary Figure 8**
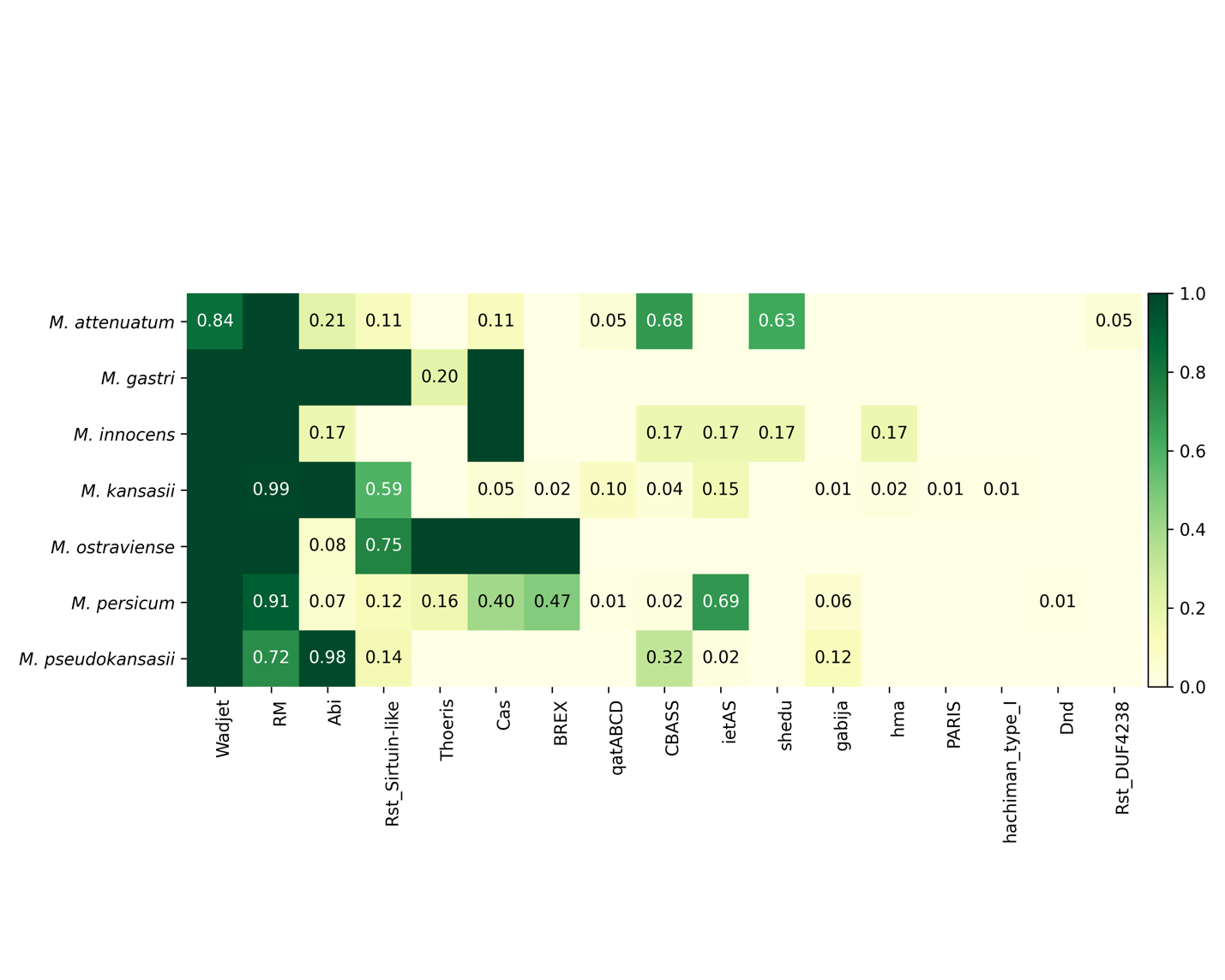

**Supplementary Figure 8. MKC defense systems.** The heatmap shows defense systems (horizontal axis) found in MKC species (vertical axis) and their proportion per species. Cells with suppressed values, indicate systems either present (dark colors) or absent (light colors) in all genomes of a given species. Numbers in cells specify the proportion of systems found only in part of genomes of a given species, with light colors representing low proportions and darker colors high proportions.

**Supplementary Figure 9**

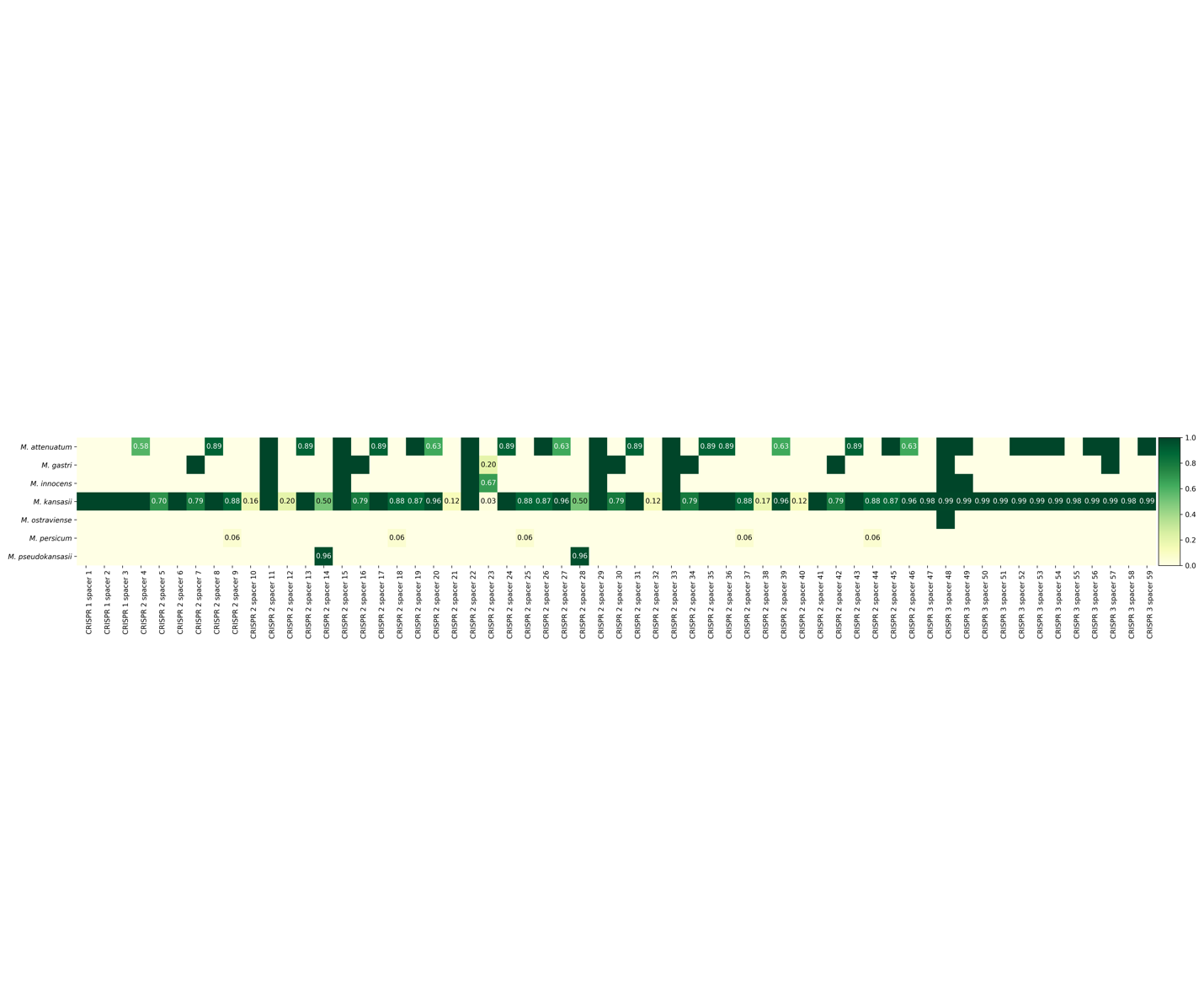

**Supplementary Figure 9**. ***M. kansasii* CRISPR loci.** The presence or absence of CRISPR spacers (horizontal) among MKC species (vertical) is represented in a heatmap. The 59 spacers belong to three CRISPR loci found in genome of the *M. kansasii* ATCC 12478 strain. Spacers either totally present (dark colors) or totally absent (light colors) in all genomes of a given species are indicated in cells with suppressed values. Alternatively, the proportion of spacers found partially in each species is explicit indicated in the respective cells, as well represented by color degree, with light for low and darker for high proportions.

**Supplementary Figure** 10

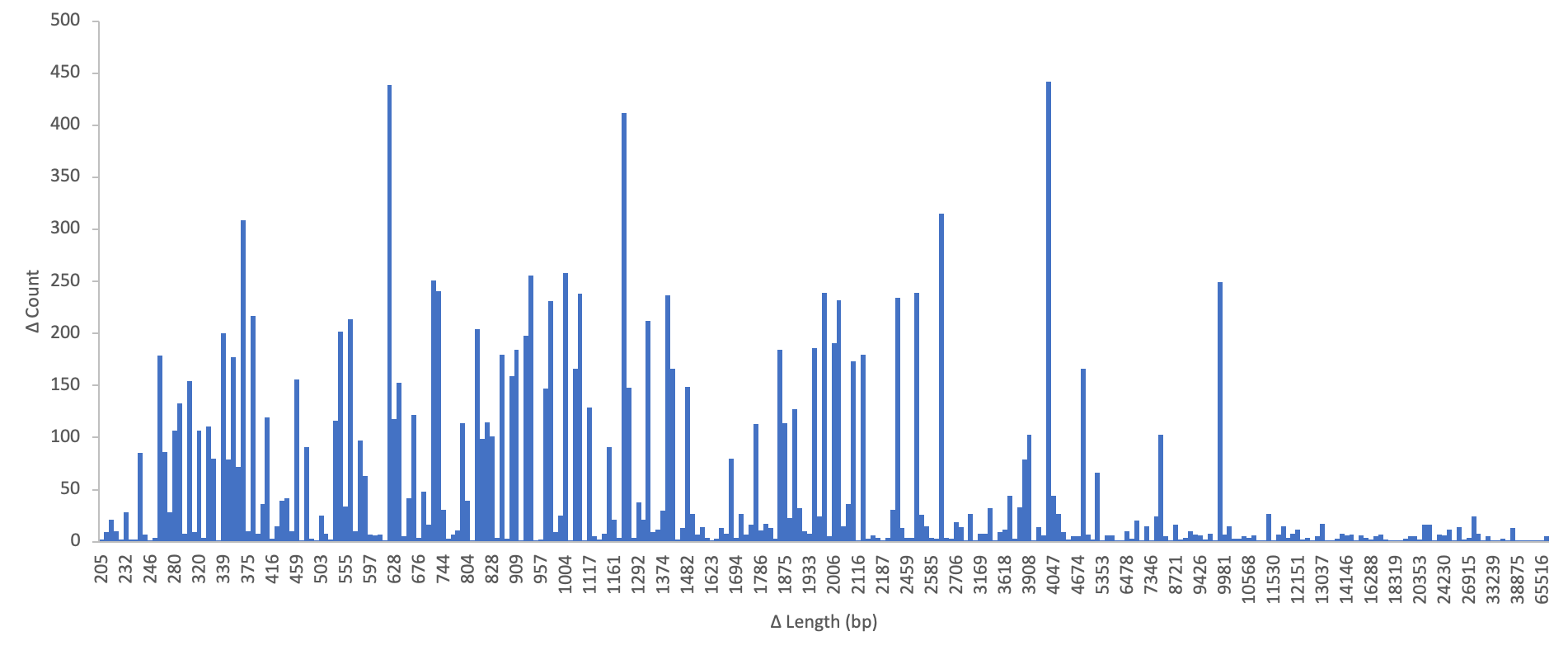

**Supplementary Figure 10. Characteristics of regions of difference in *M. kansasii* genomes.** Size distribution of regions of difference among *M. kansasii* genomes.

**Supplementary Figure 11**

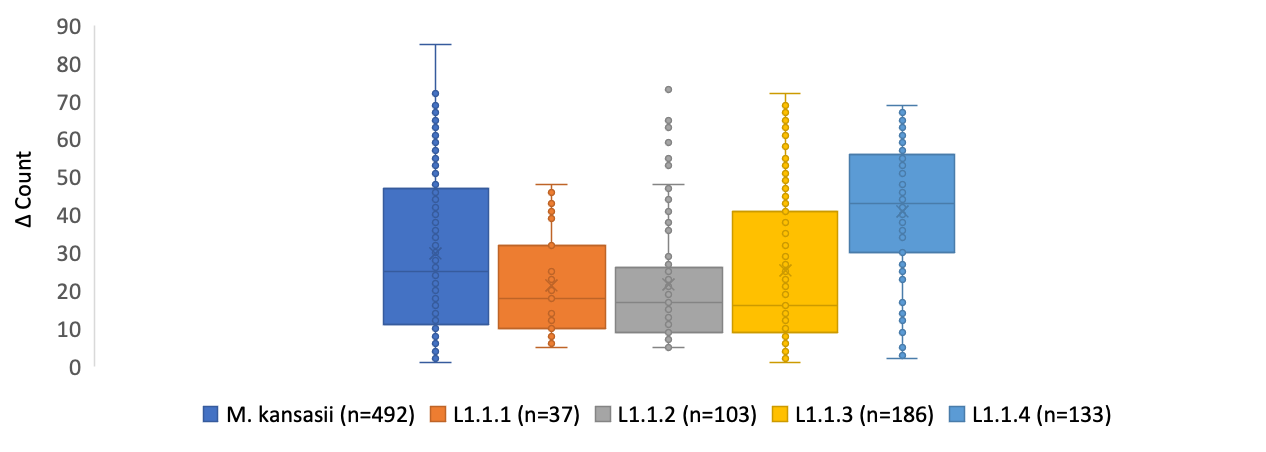

**Supplementary Figure 11. Distribution of regions of difference by *M. kansasii* sub-lineage.** Regions of difference per genome distribution among genomes of the *M. kansasii* sub-lineages. Each point represents an individual sample. The number of regions per genome is indicated in the y axis. Boxes represent the interquartile range with 50% of the values. The median is indicated by a line across the box and the mean by a “X”.
