## Supplementary figures and images for "Phylogenomic and genomic analysis reveals unique and shared genetic signatures of *Mycobacterium kansasii* complex species"

### Supplemental Data 3

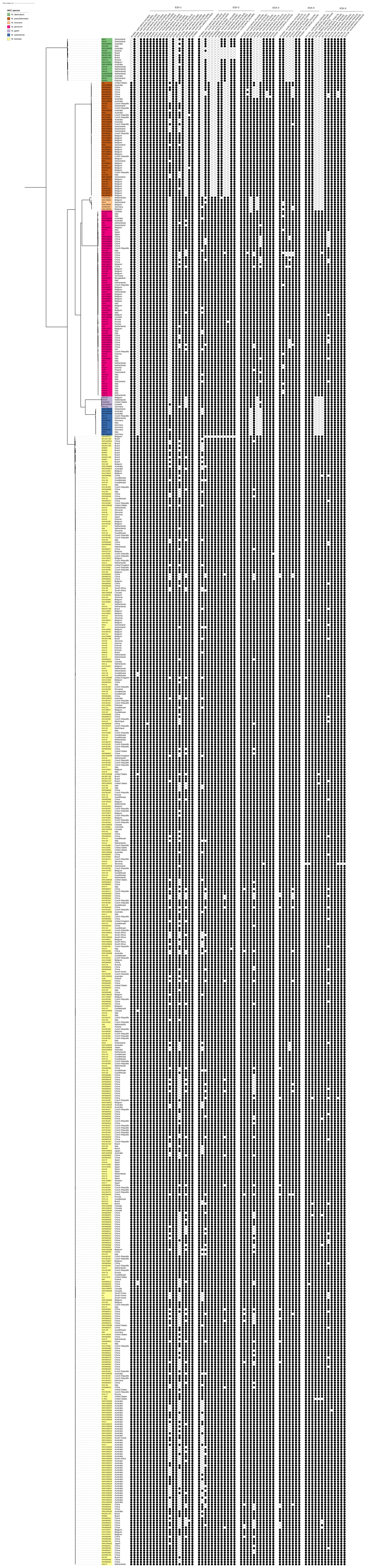

### Supplemental Data 4

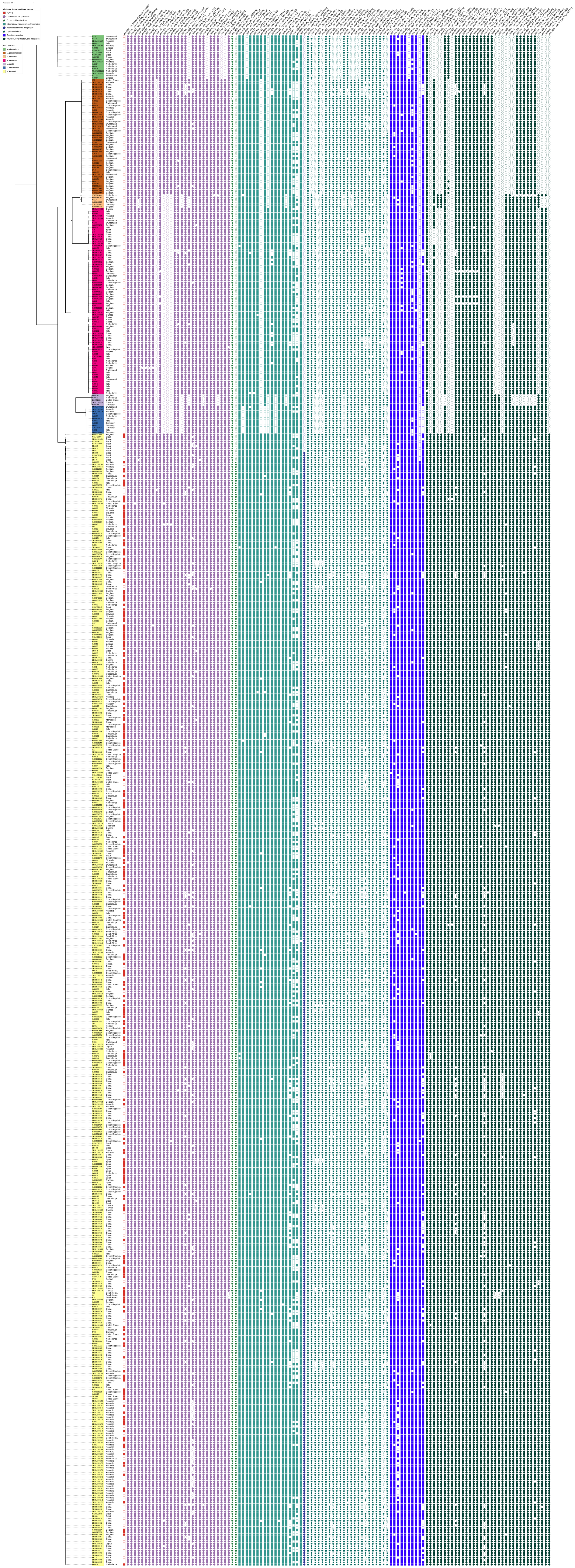
